# VadK, a Non-Canonical Kinase that Regulates the Methylcitrate Cycle and is Essential for *Mycobacterium tuberculosis* Virulence

**DOI:** 10.1101/2025.10.30.685616

**Authors:** Jordan Pascoe, Jane Newcombe, Jessica Mendoza, Shalini Birua, Tom A Mendum, Kushi Anand, Albel Singh, Amitava Sinha, Kadamba Papavinasasundaram, Apoorva Bhatt, Gerald Larrouy-Maumus, Amit Singh, Celia W Goulding, Dany JV Beste

## Abstract

The evolution of new enzymatic functions is constrained and guided by the architecture of an organism’s metabolic and regulatory networks as well as by environmental constraints. Here, we identify a previously uncharacterized kinase that has evolved from pyruvate phosphate dikinase (PPDK). Through biochemical and systems-level analyses, we show that this enzyme, encoded by Rv1127c in *Mycobacterium tuberculosis* (Mtb), has diverged from its ancestral role in central carbon metabolism to function as a histidine kinase in pathogenic mycobacteria and related species. We designate this enzyme Virulence Associated DiKinase (VadK), reflecting its ability to autophosphorylate and role in coordinating metabolism and virulence. VadK is essential for the utilization of exogenous carbon sources critical for survival within the host and is required for Mtb pathogenicity in murine models of tuberculosis. Furthermore, VadK interacts with key enzymes of the methylcitrate cycle, and ^13^C-metabolic flux analysis indicates that it fine-tunes flux through this pathway, with elevated flux proving growth limiting. Together, these findings identify VadK as a previously unrecognized regulatory kinase that integrates metabolic control with virulence in Mtb, revealing a new facet of metabolic regulation in bacterial pathogenesis and a potential target for therapeutic intervention.

## Introduction

*Mycobacterium tuberculosis* (*Mtb*), the causative agent of human tuberculosis (TB) remains a formidable human pathogen which despite decades of research and widely available treatments remains a leading cause of death globally (World Health, 2024). Although the anti-TB drug pipeline is much healthier than it has been for decades, there remains an urgent need to develop shorter, less toxic treatments that can target antibiotic resistant strains of *Mtb*. Central carbon metabolism (CCM) has been identified as a target for anti-TB drugs, an approach that has been validated by the clinical approval of bedaquiline, an ATP synthase inhibitor with high bactericidal potency (Lakshmanan & Xavier, 2013). However, CCM likely harbours many other unidentified targets with critical roles in *Mtb* survival.

Our work on Mtb metabolism previously identified that enzymes of the anaplerotic (ANA) node of CCM have unique and essential roles in controlling the flux of carbon through Mtb (Basu *et al*, 2018; Burley *et al*, 2021). The Mtb ANA node consists of the enzymes pyruvate carboxylase (PCA), phosphoenolpyruvate (PEP) carboxykinase (PCK), malic enzyme (MEZ), and Rv1127c that is annotated as pyruvate phosphate dikinase (PPDK). Whilst humans have orthologues of the enzymes PCK, MEZ and PCA the absence of PPDK in vertebrates identifies this enzyme as a promising anti-microbial drug target.

By measuring the metabolic flux profile of *Mtb* in a macrophage model of TB we discovered that in the absence of Rv1127c metabolism was dysregulated, and its growth inhibited (Basu *et al*., 2018). We also showed that Rv1127c was essential for Mtb growth in media containing cholesterol, or its breakdown product propionate, even in the presence of an additional growth permissive carbon source (Basu *et al*., 2018). Furthermore, we showed that Δ*rv1127c* is more susceptible to bedaquiline (BDQ) than wild-type (WT) Mtb (Mackenzie *et al*, 2020), and therefore drugs targeting Rv1127c may synergise with BDQ, which is the backbone of the oral, shortened treatments for antibiotic resistant TB.

In many other organisms PPDK is a critical enzyme. In bacteria and protozoa it has roles in glycolysis and ATP production via substrate level phosphorylation (Koendjbiharie *et al*, 2020), whilst in C4-plants it is critical for carbon cycling during photosynthesis (Om *et al*, 2022). Structurally PPDK is an ATP-grasp protein composed of three-distinct domains; an N-terminal ATP binding domain, a central kinase domain (containing a catalytic histidine) and a C-terminal pyruvate/PEP binding domain. PPDK can catalyse a reversable three-step reaction sequence at two separate active sites that interconverts ATP, phosphate and pyruvate to phosphoenolpyruvate (PEP), pyrophosphate and AMP (Fig S1A).

In contrast to its roles in other organisms, we have previously demonstrated using ^13^C-Metabolic Flux Analysis that in Mtb there was no flux through the PPDK reaction (Borah *et al*, 2021). Furthermore, *rv1127c* is truncated, encoding only the PPDK N-terminal ATP binding domain and the central catalytic histidine kinase domain but not the pyruvate/PEP binding domain (Fig S1B). Within the Mtb genome there were no genes encoding this domain that could form a complex with Rv1127c and so restore its canonical function (Borah *et al*., 2021). We therefore conclude that Rv1127c cannot function as a canonical PPDK.

Although *rv1127c* is not a functional PPDK, high throughput mutagenesis screens conducted by our group and others predicted that *rv1127c* is essential under a variety of conditions, including in a chemostat model (Beste *et al*, 2009), cholesterol media (Griffin *et al*, 2012), and murine models of TB (Meade *et al*, 2023; Smith *et al*, 2022).

Functionally related genes are frequently located near each other in the genome of prokaryotes. The genomic loci of *rv1127c* is adjacent to genes that encode essential enzymes of the methylcitrate cycle (MCC) (methylcitrate synthase (MCS, Rv1131c) and methylcitrate dehydratase (MCD, Rv1130c) and their transcriptional regulator (PrpR, Rv1129c) (Fig S1C) that like Rv1127c are essential for the metabolism of cholesterol and propionate (Griffin *et al*., 2012) suggesting that Rv1127c may perform an unexpected role in the MCC. The MCC is one of three approaches that Mtb can use to metabolise the potentially toxic propionyl-CoA that is derived from the catabolism of host-derived sterols, odd-chain fatty acids and branched-chain amino acids, substrates that are present and utilised in vivo (Lee *et al*, 2013). The MCC metabolises propionyl-CoA to pyruvate via the potentially toxic intermediate’s, 2-methylcitrate and 2-methylisocitrate (Russell *et al*, 2010). In the presence of vitamin B_12_ the MCC is transcriptionally repressed (Pawełczyk *et al*, 2021) and Mtb uses the alternative B_12_-dependent methylmalonyl pathway to metabolise propionyl-CoA to succinyl-CoA. Mtb can also incorporate propionyl-CoA into its cell wall including the virulence associated lipids phthiocerol dimycocerosates (PDIM) and the trehalose ester family (sulfolipid-1 (SL-1), diacyltrehalose, triacyltrehalose, and polyacyltrehalose) which are synthesised from malonyl-CoA (produced from acetyl Co-A) and methylmalonyl-CoA (produced from propionyl-CoA) (Lee *et al*., 2013). As these pathways are important for Mtb virulence and to prevent the buildup of potentially toxic metabolites, it is vital to Mtb that these pathways are rigorously regulated.

We previously demonstrated that growth attenuation of Δ*rv1127c* in propionate is rescued by the addition of either B_12_, or acetate (Basu *et al*., 2018), data that phenocopies Mtb mutants lacking MCC enzymes (Lee *et al*., 2013), thus supporting an unexpected role for Rv1127c in the MCC (Borah *et al*., 2021).

Here we demonstrate that deletion of *rv1127c* results in dysregulated CCM, disruption in redox balance, and in bioenergetics when grown on propionate-containing, or yielding medium, and reduced survival and virulence of Mtb in two different murine models of tuberculosis. Our results confirm the importance of this protein in the life cycle of Mtb and therefore we herein rename this protein <u>V</u>irulence <u>A</u>ssociated <u>Dik</u>inase (VadK). We confirm that despite being truncated Rv1127c is a functional histidine kinase that fine-tunes the MCC by interacting with the enzymes of the MCC pathway. The discovery of a unique, non-canonical histidine phosphotransfer system that has evolved from the CCM enzyme PPDK in pathogenic Mycobacteria and other related genera enhances our understanding of the regulation of metabolism and has implications for understanding the role of the MCC in Mtb growth and virulence.

## Methods

### Bacterial strains

Frozen stocks of *Mtb* (H37Rv) and mutant strains *ΔvadK* and Δ*vadK:vadK* are previously described (Basu *et al*., 2018). For co-immunoprecipitation the overexpression vector pKP1153 was constructed by cloning C-terminally FLAG-tagged Rv1127c under the control of the constitutive mycobacterial Ptb38 promoter into pDE43-MEH using a Gateway cloning approach (Ehrt *et al*, 2005; Kim *et al*, 2011). Site directed mutagenesis of pMV306ppdk (Basu *et al*., 2018) by Genescript generated 3 different alleles of *vadK: vadK*^His422Ala^, vadK^His422Asp^ and vadK^Arg106Lys^. All plasmids were transformed by electroporation into WT or *ΔvadK* and selected on 7H11 plates supplemented with 25 μg ml^-1^ kanamycin or 50 μg ml^-1^ hygromycin, as appropriate. (Basu *et al*., 2018). PCR was used to confirm the presence of the correct gene.

### Growth conditions

*Mtb* strains were cultivated using Middlebrook 7H11 agar containing 5% (v/v) oleic acid/albumin/dextrose/catalase (OADC) enrichment medium supplement (BD) and 0.5% (v/v) glycerol. Liquid cultures were grown in standard Middlebrook 7H9 broth containing 0.2% (v/v) glycerol, 0.2% (v/v) Tween 80 or Tyloxapol, and 5% (v/v) ADC or Rosins minimal media (Beste *et al*, 2005). For the methylcitrate synthase assay, strains were grown in 7H9 media containing 0.2% (v/v) glycerol, 0.2% (v/v) Tyloxapol, and 5% (v/v) ADC and 10- or 20-mm sodium propionate. Vitamin B12 (10 μg ml^−1^) was added when indicated.

For the sole carbon source experiments Roisin’s minimal media (Beste *et al*., 2005) containing either glycerol (0.5%), pyruvate (0.165%) or cholesterol (0.154%) and 0.2% tyloxapol. Cultures were grown until mid-log phase (OD_600nm_ = 0.6-0.8) in standard 7H9, washed once with PBS, and then resuspended to a starting OD_600nm_ of 0.01). Cell growth was monitored daily by OD and/or CFU measurements. When selection was required, kanamycin at 20 μg ml^−1^, hygromycin at 50 μg ml^−1^ were added to the culture medium.

### Animal Ethics and Biosafety Approvals

All animal experiments were conducted in strict accordance with the guidelines of the Committee for the Purpose of Control and Supervision of Experiments on Animals (CPCSEA), Government of India, and were approved by the Institutional Animal Ethics Committee (IAEC: CAF/Ethics/780/2020) and the Institutional Biosafety Level-3 (BSL-3) Committee.

### Mouse Infection Models

For Infection studies, 6–8-week-old both male and female C3HeB/FeJ mice and female BALB/c mice (n=7-8 per group) were infected by aerosol using Glas-Col Inhalation Exposure System to deliver approximately 50 bacilli per mouse with WT, knockout and complemented Mtb strains as indicated (Mtb H37Rv, MtbΔvadk and MtbΔvadk:vadk). Mice were housed inside the BSL-3 facility and euthanized at day 28 and day 56 post-infection. Lungs and spleens were aseptically harvested and homogenized in sterile PBS for bacillary load analysis, tissue histopathology, and pathological scoring, as described [PMID: 34392160]. Serial dilutions of tissue homogenates were plated on Middlebrook 7H11 agar plates supplemented with OADC enrichment and PANTA antibiotic mixture. Plates were incubated at 37 °C, and CFUs were enumerated after 4 weeks of incubation

### Histopathological Analysis

Lung tissues were fixed in 10% formaldehyde, embedded in paraffin, and sectioned for histological evaluation. Sections were stained with Hematoxylin and eosin (H&E) for pathological examination. Disease severity was assessed based on granuloma scoring and representative histopathological images were captured for each infection group.

### In vitro VadK expression and purification

To improve the solubility of recombinant VadK we expressed a slightly truncated version of the protein lacking 17 N-terminal residues that are predicted to be disordered according to AlphaFold3 (Abramson *et al*, 2024) predictions (Fig S2A). We also found that co-expressing VadK with its operonic partner Rv1126c improved its solubility. The synthetic Mtb Rv1127c gene sequence, codon-optimized for *Escherichia coli*, was cloned into the pET28a-TEV vector (GenScript) to produce an N-terminal TEV-cleavable 6x His-tagged VadK. The neighbouring gene, Rv1126c gene sequence was codon-optimized for *E. coli* and cloned into pET22b (Novagen) to produce Rv1126c without a HisTag. Successful cloning was verified by DNA sequencing (GeneWiz from Azenta Life Sciences). Similarly, VadK^His422Ala^ was constructed in pET28a-thrombin vector. Site-directed mutagenesis was performed using Pfu Ultra DNA Polymerase (Agilent) and Rv1127c H422A-forward (5’- TGGTGCGGCCTCCGCAGCAGCGGTGGTGTCACGTGAACTA) and Rv1127c H422A-reverse (sequence: 5’- TAGTTCACGTGACACCACCGCTGCTGCGGAGGCCGCACCA) primers (Azenta). The VadK^His422Ala^ mutant was confirmed by DNA sequencing (GeneWiz from Azenta Life Sciences).

pET28a-TEV-VadK-HisTag and pET22b-Rv1126c-noTag, were co-transformed into *E. coli* BL21-Gold (DE3) T7 Express cells (New England Biolabs Inc) and were grown in LB media containing 100 μgml^-1^ ampicillin and 50 µgml^-1^ kanamycin and protein expression was induced by 1 mM IPTG. Cultures were lysed by sonication on ice (45 % amplitude, 15 seconds on and 45 seconds off for 15 cycles) in Buffer A (50 mM Tris pH 7.4, 350 mM NaCl, 10 mM imidazole), 100 µM PMSF and 0.5 mg ml^-1^ lysozyme). Proteins were loaded onto a HisTrap FF column (5 mL, GE Healthcare) using an AKTA Start FPLC and washed extensively with Buffer A containing 2M NaCl to seperate Rv1126c from Rv1127c. Bound protein was eluted from the column using an imidazole gradient (10mM to 0.5 mM) in Buffer A. Rv1127c was pooled, concentrated, and further purified using a Superdex 200 Increase 10/300 GL column (Cytiva) equilibrated with Buffer B (50 mM Tris pH 7.4, 150mM NaCl buffer, 10% glycerol) to yield pure fractions of VadK-HisTag (Fig S2B). The expression and purification method for Rv1127c^H422A^ mutant was carried out in an identical fashion.

TEV at a ratio of 10:1 VadK:TEV was used to remove the His-tag from VadK in Buffer B with 2 mM DDT at 4 °C. Cleaved VadK was recovered using NiNTA agarose beads (ThermoFisherScientific) and further purified using a Superdex 200 Increase 10/300 GL column (Cytiva) equilibrated with Buffer B. Pure fractions were pooled and concentrated.

### VadK autophosphorylation assays

Reaction mixtures of 25 μM VadK, 2mM ATP and 2mM MgCl_2_ in Buffer B were incubated for 2 hours at room temperature. Samples were analysed by mass spectrometry (MS) using LC-MS/MS (ACQUITY UPLC H-class system, Xevo G2-XS QTof, Waters). Phosphorylation was observed by reverse-phase chromatography at 45 °C using a C4 column (Protein BEH C4 Column, 300Å, 1.7 μm, 2.1 mm X 50 mm, Waters) and a 5-minute gradient of Buffer C (0.1% formic acid in water), Buffer D (100% acetonitrile from 0% Buffer D to 90% Buffer D with a flow rate of 0.3 mL/min). The Xevo Z-spray source was run with a capillary voltage of 300 V, and a cone voltage of 40 V (NaCsI calibration, Leu-enkephalin lock-mass). N_2_ was used as the desolvation gas at 350 °C and a flow rate of 800 L/hour and data acquisition was done at alternating collision energy with low energy at 0 V and high collision energy ramp from 15-45 V. Data was acquired in continuum with 0.5 second scans across a mass range of 400-4000 Da. Data was deconvoluted using proprietary Water MaxEnt1 algorithm.

### Circular dichroism (CD) of Mtb VadK and its mutant

Far UV CD spectra were collected at 25°C using a Jasco J-810 spectropolarimeter with a Digital Integration Time of 1 second and a bandwidth of 1 nm, using 0.1 cm cuvettes with VadK and its mutant (5 µM) in 5 mM Tris pH 7.4, 35 mM NaCl, 1% glycerol. Spectra were read from 260-190nm at 100 nm/min for a total of 10 accumulations. The BeStSel tool (https://bestsel.elte.hu/index.php) was used to quantify secondary structural elements.

### VadK sequencing by tandem MS to confirm the catalytic histidine

Reaction mixture (25 μM VadK, 2 mM ATP, 2 mM MgCl_2_, in Buffer B) was incubated for 2 hours at room temperature. Samples of apo and phosphorylated VadK (4 µM) and DTT (20 mM) in 50 mM ammonium bicarbonate (ABC) buffer, pH 7.4 were boiled at 80°C for 20 minutes. After cooling, iodoacetamide (15 mM) was added to the samples and incubated in the dark at room temperature for 1 hour, followed by the addition of Trypsin Gold (400 nM, Promega) f or an overnight digestion at 37°C. Samples were analysed using liquid chromatography tandem MS (LC-MS/MS, ACQUITY UPLC H-class system, Xevo G2-XS QTof, Waters) by reverse-phase chromatography at 45 °C using a C4 column (Protein BEH C4 Column, 300Å, 1.7 μm, 2.1 mm X 50 mm, Waters) and a 5-minute gradient of Buffer C and Buffer D with a flow rate of 0.3 mL min-1. The Xevo Z-spray source was run with a capillary voltage of 300 V, and a cone voltage of 40 V (NaCsI calibration, Leu-enkephalin lock-mass). N_2_ was used as the desolvation gas at 350 °C and a flow rate of 800 L/hour and data acquisition was carried out at alternating collision energy with low energy at 0 V and high collision energy ramp from 15-45 V. Data was acquired in continuum with 0.5 second scans across a mass range of 400-4000 Da. Waters proprietary software BioPharma Lynx was used to identify phosphorylated peptide fragment, intensities, and control b/y ions. Finally, the b/y ions were then further identified in raw Xevo MS/MS data.

### Co-Immunoprecipitation of FLAG-tagged VadK

We followed the methodology described here (de Miranda *et al*, 2023). Briefly, 50 ml cultures of Mtb were grown in 7H9 media with 0.2% glycerol and Tyloxapol with and without 10 mM sodium propionate until late log phase (OD_600_ = 0.8 – 1). Cell pellets were resuspended in lysis buffer (50 mM Tris pH7.4, 350 mM NaCl,10% glycerol, Complete protease inhibitor (Roche) by bead-beating with lysis matrix B using a Hybaid Fast Prep at speed (4.0) for 20 seconds with careful chilling between each round. Lysates were incubated with 1% n-Dodecyl-B-D-maltoside (Avanti) for 4 h at 4°C before centrifugation and then filtered twice through 0.22 μm Spin-X column filters (CoStar). Anti Flag M2 affinity matrix (40 µl) (Sigma) that had been pre-washed in TBS (Tris-Buffered Saline, 50 mM Tris-HCl pH 7.4, 150 mM NaCl) was added to the lysate and incubated overnight rotating at 4°C. The resin was carefully washed 5 times with TBS before being resuspended in NuPAGE LDS loading dye and heated at 95°C for 5 m. Proteins were electrophoretically separated on a 10% Novex Bis-Tris SDS -PAGE gel (Invitrogen) with MES buffer and the entire gel slice was sent for MS analysis.

Each gel slice was subjected to in-gel tryptic digestion using a DigestPro automated digestion unit (Intavis Ltd.). The resulting peptides were fractionated using an Ultimate 3000 nano-LC system in line with an Orbitrap Fusion Lumos mass spectrometer (Thermo Scientific). In brief, peptides in 1% (vol/vol) formic acid were injected onto an Acclaim PepMap C18 nano-trap column (Thermo Scientific). After washing with 0.5% (vol/vol) acetonitrile 0.1% (vol/vol) formic acid peptides were resolved on a 250 mm × 75 μm Acclaim PepMap C18 reverse phase analytical column (Thermo Scientific) over a 150 min organic gradient, using 7 gradient segments (1-6% solvent B over 1min., 6-15% B over 58 min., 15-32% B over 58min., 32-40% B over 5min., 40-90%B over 1min., held at 90%B for 6min and then reduced to 1%B over 1min.) with a flow rate of 300 nl min−1. Solvent A (0.1% formic acid), and Solvent B (80% acetonitrile in 0.1% formic acid). Peptides were ionized by nano-electrospray ionization at 2.2 kV using a stainless-steel emitter with an internal diameter of 30 μm (Thermo Scientific) and a capillary temperature of 300°C.

All spectra were acquired using an Orbitrap Fusion Lumos mass spectrometer controlled by Xcalibur 3.0 software (Thermo Scientific) and operated in data-dependent acquisition mode. FTMS1 spectra were collected at a resolution of 120 000 over a scan range (m/z) of 350-1550, with an automatic gain control (AGC) target of 4E5 and a max injection time of 50ms. Precursors were filtered according to charge state (to include charge states 2-7), with monoisotopic peak determination set to peptide and using an intensity threshold of 1E3. Previously interrogated precursors were excluded using a dynamic window (40s +/-10ppm). The MS2 precursors were isolated with a quadrupole isolation window of 0.7m/z. ITMS2 spectra were collected with an AGC target of 2E4, max injection time of 35ms and HCD collision energy of 30%. The raw data files were processed using Proteome Discoverer software v1.4 (Thermo Scientific) and searched against the UniProt *Mycobacterium tuberculosis* (strain ATCC 25618 H37Rv) [83332] database (downloaded July 2022; 3993 sequences) and the XX sequence using the SEQUEST HT algorithm. Peptide precursor mass tolerance was set at 10ppm, and MS/MS tolerance was set at 0.6Da. Search criteria included oxidation of methionine (+15.995Da) and phosphorylation of histidine (+79.966Da) as variable modifications and carbamidomethylation of cysteine (+57.021Da) as a fixed modification. Searches were performed with full tryptic digestion, and a maximum of two missed cleavages were allowed. The reverse database search option was enabled, and all data was filtered to satisfy false discovery rate (FDR) of 1%.

### RNA-sequencing

*Mtb* strains were grown in standard 7H9 until mid-log phase (OD_600_= 0.6 - 0.8) before being spiked with 20 mM sodium propionate. After 4 h transcription was stopped by the addition of 4 volumes GTC buffer (5M guanidine isothiocyanate, 0.5% sodium L-lauryl sarcosine, 25mM tri-sodium citrate pH7. 0.1M 2-mercaptoethanol). RNA was extracted as described (Beste *et al*, 2007). RNA was treated with DNAse twice and purifed (RNAeasy kit (NEB)). RNA sequencing and analysis was performed by Genewiz (Azenta). RNA samples were quantified using Qubit 4.0 Fluorometer (Life Technologies, Carlsbad, CA, USA) and RNA integrity was checked with RNA Kit on TapeStation (Agilent Technologies, Palo Alto, CA, USA). ERCC RNA Spike-In Mix (Cat: #4456740) from ThermoFisher Scientific, was added to normalized total RNA prior to library preparation following manufacturer’s protocol. RNA depletion was performed using NEBNext rRNA Depletion Kit (Bacteria). RNA sequencing library preparation used NEBNext Ultra RNA Library Prep Kit for Illumina by following the manufacturer’s recommendations (NEB, Ipswich, MA, USA). Briefly, enriched RNAs were fragmented according to manufacturer’s instruction. First strand and second strand cDNA were subsequently synthesized. cDNA fragments were end repaired and adenylated at 3’ends, and universal adapter was ligated to cDNA fragments, followed by index addition and library enrichment with limited cycle PCR. Sequencing libraries were validated using NGS Kit on the Agilent 5300 Fragment Analyzer (Agilent Technologies, Palo Alto, CA, USA), and quantified by using Qubit 4.0 Fluorometer (Invitrogen, Carlsbad, CA).

The sequencing libraries were multiplexed and loaded on the flowcell on the Illumina NovaSeq X Plus instrument according to manufacturer’s instructions. The samples were sequenced using a 2x150 Pair-End (PE) configuration. Image analysis and base calling were conducted by the NovaSeq Control Software on the NovaSeq instrument. Raw sequence data (.bcl files) generated from Illumina NovaSeq was converted into fastq files and de-multiplexed using Illumina bcl2fastq program. One mismatch was allowed for index sequence identification.

Sequence reads were trimmed to remove possible adapter sequences and nucleotides with poor quality using Trimmomatic v.0.36. The trimmed reads were then mapped to the Mtb H37Rv reference genome using the Bowtie2 aligner v.2.2.6 to generate BAM files. Unique gene hit counts were calculated by using featureCounts from the Subread package v.1.5.2. The hit counts were summarized and reported using the $gene_feature feature in the annotation file. Only unique reads that fell within gene regions were counted.

After extraction of gene hit counts, the gene hit counts table was used for downstream differential expression analysis. Using DESeq2, a comparison of gene expression was performed. The Wald test was used to generate p-values and log2 fold changes. Genes with < 0.01 padj value and absolute log2 fold change > 1.2 were called as differentially expressed genes for each comparison.

### 13C-isotopologue Analysis

Mtb strains were grown in standard 7H9 media to late log phase (OD_600_ = 0.8-1) before being washed and resuspended in ^13^C labelled media and incubated for 48 h. Using [U-13C3]-glycerol (99%), 20 mM [U-13C3]-pyruvate (99%) or 0.154% [3, 4- 13C2] cholesterol for 48 h. Metabolites were quenched and extracted using a slight modification of the protocol described (Thomson *et al*, 2022). The bacteria were metabolically quenched by plunging into acetonitrile/methanol/H2O (2:2:1) precooled to -80 °C. Metabolites were extracted by bead beating in the FastPrep at 6.5 for 20 s x 3 with careful cooling in between. Soluble extracts were filtered twice through 0.22 μm Spin-X column filters (CoStar) and then stored at −80°C until analysis.

The data were acquired with an Agilent 1290 Infinity II UHPLC coupled to a 6545 LC/Q-TOF system. Chromatographic separation was performed with an Agilent InfinityLab Poroshell 120 HILIC-Z (2.1 × 100 mm, 2.7 μm (p/n 675775-924)) column. The HILIC-Z methodology was optimized for polar acidic metabolites. Column compartment was set at 50°C. For easy and consistent mobile-phase preparation, a concentrated 10 × solution consisting of 100 mM ammonium acetate (pH 9.0) in water was prepared to produce mobile phases A and B. Mobile phase A consisted of 10 mM ammonium acetate in water (pH 9) with a 5 μM Agilent InfinityLab deactivator additive (p/n 5191-4506), and mobile phase B consisted of 10 mM ammonium acetate (pH 9) in 10:90 (v:v) water/acetonitrile with a 5 μM Agilent InfinityLab deactivator additive (p/n 5191-4506). The following gradient was applied at a flow rate of 0.25 ml/min: 0 min, 96% B; 2 min, 96% B; 5.5 min, 88% B; 8.5 min, 88% B; 9 min, 86% B; 14 min, 86% B; 17 min, 82% B; 23 min, 65% B; 24 min, 65% B; 24.5 min, 96% B; 26 min, 96% B and 3-min of re-equilibration at 96% B. Accurate MS was performed using an Agilent Accurate Mass 6545 QTOF apparatus. Dynamic mass axis calibration was achieved by continuous infusion after the chromatography of a reference mass solution using an isocratic pump connected to an electrospray ionization source operated in negative-ion mode. The following parameters were used: gas temperature, 225°C; drying gas, 13 l min-1; sheath gas temperature, 350 °C; nebulizer pressure, 35 psi; sheath gas flow, 12 l min-1; capillary voltage, 3,500 V; nozzle voltage, 0 V; fragmentor voltage, 125 V; skimmer 45V and octupole 1 RF voltage, 750V. The data were collected in centroid 4 GHz (extended dynamic range) mode.

### LC-MS metabolomics data analysis

Data analysis was performed using the Agilent MassHunter Qualitative (v10), Quantitative Analysis and Profinder Softwares. Metabolite identification was based on mass-retention times and isotope distribution patterns. Metabolites were quantified using area under the curve (AUC) normalized to protein concentration determined using a standard Bradford assay.

### Determination of ATP levels

ATP was measured using the luciferase-based ATP determination kit (Thermofisher, US; A22066) according to the manufacturer’s instructions. Strains of Mtb were grown in standard 7H9 media with or without 10 mM propionate for 14 days. Samples were harvested after 7- and 14-days incubation and extracted in lysis buffer (50 mM Tris pH7.4, 150 mM NaCl_2_ and 0.01% Tween80) by bead-beating with 0.1 mm zirconia-silica beads (Sigma) in the FastPrep at 6.5 for 3 × 45 s with careful cooling between bursts.

### Analysis of mycobacterial Lipids

Mtb strains were in standard Middlebrook 7H9 media for 10 days before spiking with 20 mM sodium propionate and a further 5-days incubation. Cell pellets were washed with PBS and autoclaved before extraction. Extraction of Mtb lipids and Thin Layer Chromatography (TLC) analysis was carried out using protocols described by Dobson *et al* (1985) (DOBSON *et al*, 1983). Dry weights of cell pellets were used as a measure to equalize loading on TLC plates using solvent Systems A (direction 1; Petroleum ether/ethyl acetate 98:2 x 3, direction 2; Petroleum ether/acetone 98:2), System B (direction 1; Petroleum ether/acetone 92:8 x 3, direction 2; Toluene/acetone 95:5), System C (direction 1; Chloroform/methanol 96:4, direction 2; Toluene/acetone 80:20), System D, (direction 1; chloroform/methanol/water 100:14:0.8, direction 2; chloroform/acetone/methanol/water 50:60:25:3)

### Measurement using Mrx1-roGFP2 redox biosensor

The Mrx1-roGFP2 ratio was determined during *in vitro* growth of *Mtb* strains in glycerol (0.2 %) or propionate (10 mM and 20 mM) containing 7H9 medium as described in (Das *et al*, 2023). Briefly, bacterial cultures expressing Mrx1-roGFP2 were treated with 5 mM *N*-ethylmaleimide (Sigma-Aldrich, St. Louis, MO) for 5 min at room temperature followed by 4% paraformaldehyde (PFA) fixation (Himedia, Mumbai, India) for 1 hour at room temperature. Bacteria were analysed using a FACSVerse flow cytometer (BD Biosciences, San Jose, CA). The biosensor response was quantified by measuring the fluorescence ratio at a fixed emission (510 nm) on excitation at 405 and 488 nm. The data obtained were analysed with the BD FACSuite software. These ratiometric data were normalized to measurements of cells treated with 10 mM CHP (Sigma-Aldrich, St. Louis, MO), giving maximal oxidation of the biosensor, and 20 mM dithiothreitol (Sigma-Aldrich, St. Louis, MO), yielding a readout of maximal reduction of the biosensor. Ten thousand events per sample were analysed.

### OCR and ECAR measurements

To assess the metabolic activity of *Mtb* strains, the basal oxygen consumption rate (OCR) and extracellular acidification rate (ECAR) were measured using the Agilent Seahorse XFp Analyzer. Log-phase *Mtb* cultures were subjected to nutrient starvation in Middlebrook 7H9 broth lacking albumin, dextrose, and sodium chloride (ADS) or any carbon source, and supplemented with tyloxapol, a non-metabolizable detergent, to prevent bacterial clumping. Following starvation, cultures were centrifuged at 100 × g for 5 minutes to enrich for single-cell suspensions. The resulting suspensions were washed thoroughly with unbuffered 7H9 medium to remove residual nutrients. A total of 4 × 10⁶ bacterial cells per well were seeded into the wells of a Cell-Tak–coated XF cell culture microplate (Corning, Cat. No. 354240). To ensure equal seeding density across wells, CFU validation was performed by plating aliquots on Middlebrook 7H11 agar and enumerating colonies after a 4-week incubation at 37 °C. OCR and ECAR were measured using unbuffered 7H9 assay medium (pH 7.45; lacking disodium phosphate and monopotassium phosphate). The medium was supplemented with 20 mM sodium propionate and 0.2% glycerol as the sole carbon sources. Measurements were recorded over a period of approximately 60 minutes. To assess respiratory capacity, carbonyl cyanide 3-chlorophenylhydrazone (CCCP) (Sigma-Aldrich, Cat. No. C2759) was added at a final concentration of 10 μM through designated drug injection ports during the assay, as indicated in corresponding figure legends. Changes in OCR and ECAR in response to substrate addition or CCCP stimulation were calculated and expressed as pmol/min per 4 × 10⁶ cells.

### Crude Enzyme Extractions

Mtb was grown to late log phase (OD_600=_ 0.8–1.0) in standard 7H9 broth. Bacterial pellets were harvested and resuspended in the same media with or without 10 mM propionate and incubated for 1 or 4 days. Preparation of cell-free extracts was performed basically as (Muñoz-Elías *et al*, 2006). Briefly bacterial pellets were lysed in buffer (150mM NaCl_2_, 50mM Tris, 10% (v/v) glycerol, 1 mM phenylmethylsulfonyl fluoride (PMSF) at pH 7.4) by bead-beating as described for the ATP assay. Extracts were clarified by centrifugation and filtration through a 0.45µm pore SpinX filter (Corning) and the total protein concentration was determined by BCA assay. Cell-free extracts were standardised to 0.25 mg ml^-1^frozen and stored at −80°C.

### Methylcitrate synthase assay

The enzyme activity of crude extracts was performed as described by (Muñoz-Elías *et al*., 2006), with some modifications. Briefly, MCS activity was measured at 30^°^C using the reaction buffer: 50mM Tris pH 7.4, 150 mM NaCl_2_, 2 mM 5,5-dithio-bis-(2-nitrobenzoic acid (DTNB), 0.5 mM oxaloacetate (OAA), 0.3 mM propionyl-CoA. Activity was measured at spectrophotometrically at 400 nm.

### Phylogenetic analysis

PPDK sequences of typical mycobacteria and other relevant organisms were downloaded from UniProt (Consortium, 2024). The evolutionary history was inferred with MEGA X (Kumar *et al*, 2018), sequences were aligned using the MUSCLE algorithm and a phylogeny generated using the Maximum Likelihood method with the optimised Whelan and Goldman + Freq. model (Whelan & Goldman, 2001).

## Results

### VadK is essential for Mtb to cause tuberculosis

We previously demonstrated that Rv1127c (VadK) is essential for the intracellular growth of Mtb in macrophages, and this correlated with significant dysregulation of intraphagosomal bacterial metabolism, suggesting that VadK plays a critical role in bacterial survival within the host (Basu *et al*., 2018). To test this hypothesis, we used two murine models of TB, a low-dose aerosol infection of BALB/c mice and the Kramnik model (C3HeB/FeJ mice) which develop human-like granulomas (Kramnik *et al*, 2000). In accordance with our hypothesis Δ*vadK* was severely attenuated, unable to grow in the lungs or disseminate into the spleen of BALB/c mice (Fig S3A & B). Although there was a significant difference between the growth of Δ*vadK* and Δ*vadK:vadK* complementation was incomplete suggesting that there was insufficient expression of *vad*K from the chromosomally distant *att*B location where we had re-inserted *vad*K back into the genome. To eliminate the possibility of polar effects, the Δ*vadK* genome was sequenced but no additional mutations were identified as compared with the parental strain. Further, the histopathology data (Fig S3C & D) showed that whilst mice infected with Δ*vadK* had significantly less tissue damage there were no significant differences between WT and *ΔvadK:vadK* infected mice providing further evidence that the phenotype of Δ*vadK* was legitimate.

The bacterial burden in C3HeB/FeJ mice showed a similar result to the BALB/C model again demonstrating that Δ*vadK* was unable to grow in either the lungs nor disseminate to the spleen (Fig 1A & B). Complementation restored bacterial burden in the lungs and dissemination to the spleen. Consistent with this, Δ*vadK* infection caused significantly less pathology (granuloma score,15.3) as compared with the parental (granuloma score, 68.3) and the complemented strain (granuloma score, 57.7) (Fig 1C & D). In conclusion, the infection experiments showed that VadK is required for *Mtb* to cause disease in murine models of tuberculosis and thus essential for virulence.

**Figure 1.**
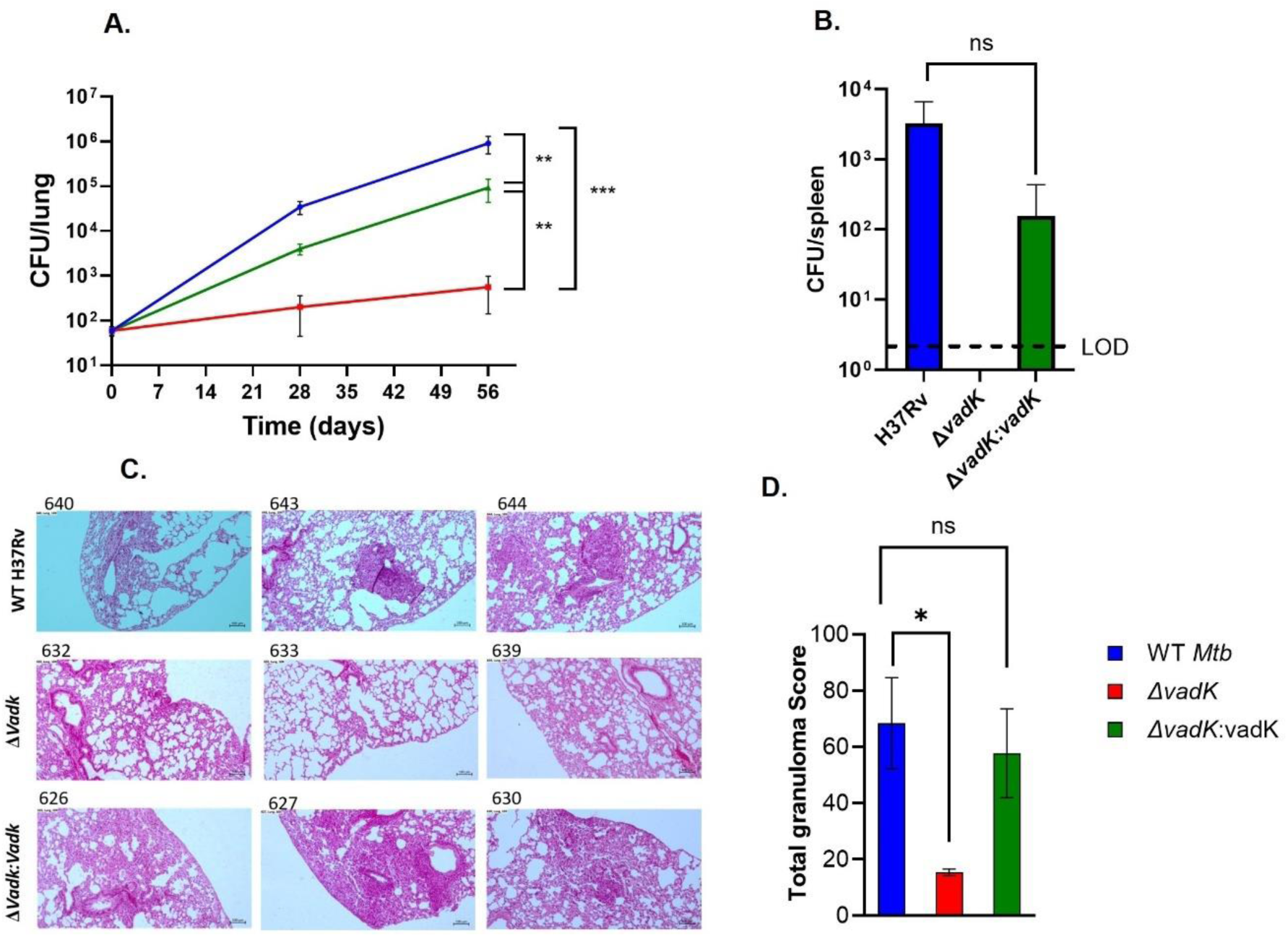
VadK is required for Mtb to cause tuberculosis in C3HeB/FeJ mice. C3HeB/FeJ mice (n=8) were aerosol infected with WT (blue), Δ*vadK* (red) and Δ*vadK:vadK* (green). Mtb survival was measured in the lungs at the indicated time points (A) and dissemination to the spleen after 56 days (B). The limit of detection (LOD) is 20 CFU’s. Lung sections were stained with Haematoxylin-and-eosin after 56 days of infection and scored blindly by a pathologist using (Kramnik & Beamer, 2016). The images show HE-stained (10 X magnification) from individual lung sections representative of three infected animals/group (C) and the scores for animals in each group with mean± standard error mean (SEM). The data depicted are means ± SEM for each group. P>0.05:NS, P<0.001:***, p<0.01:**, p<0.05:*, unpaired two-tailed t-test with Welch’s correction.

### VadK can autophosphorylate *in vitro*

VadK consists of an N-terminal ATP binding domain and a phospho-transfer histidine kinase domain. We predicted that VadK is an ATP-dependent (di)kinase that autophosphorylates its catalytic histidine. To test this, we purified recombinant VadK (Fig S2B) and tested its ability to autophosphorylate by measuring ATP-dependent VadK phosphorylation with and without Mg^2+^ (an essential co-factor of PPDK) by LC-MS (Xevo G2-XS QTof). After 2 h incubation at RT, we detected both apo-VadK at the expected mass/charge (m/z=50498) and phosphorylated-VadK with an increased mass shift of ∼80 Da in accordance with a phosphorylation event (Fig 2). Phosphorylation was dependent on the presence of both Mg^2+^ and ATP (Fig 2). These results show that VadK despite being truncated retains its kinase activity.

**Figure 2.**
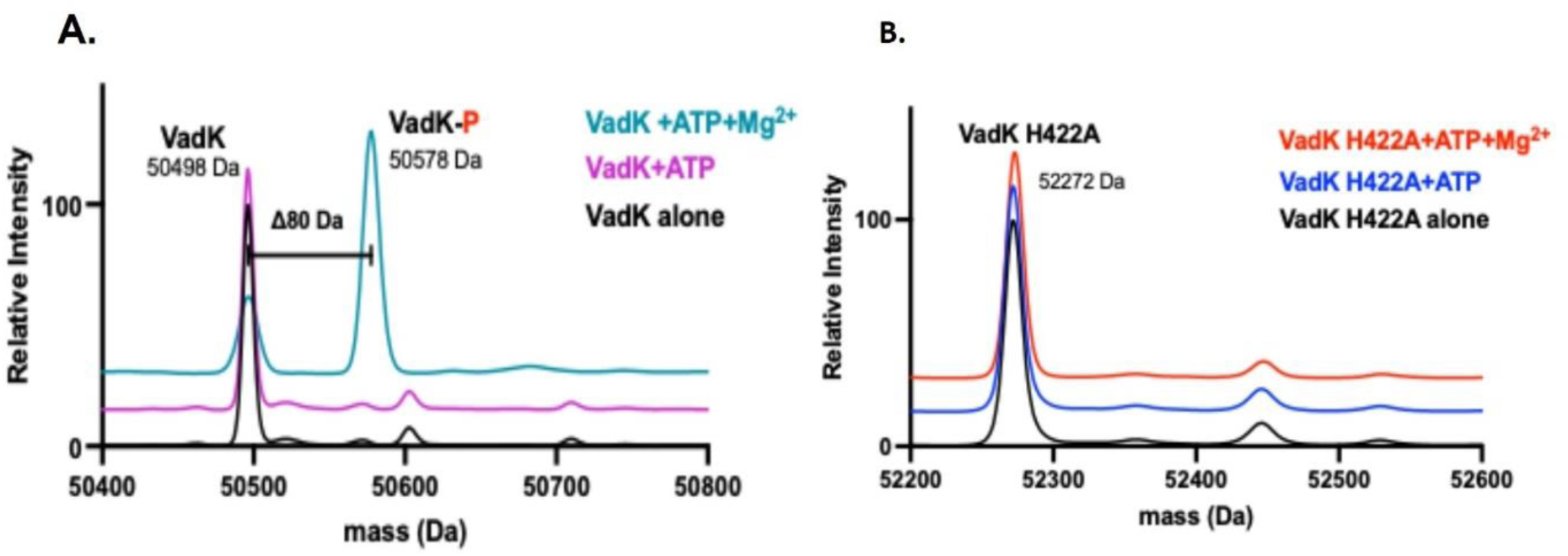
VadK-His can auto-phosphorylate. (A) MS of VadK alone (black line), with ATP and in the absence (purple line) and presence of Mg_2+_ (cyan line). A phosphorylation event (mass shift 80 Da, designated by VadK-P) only occurs for apo-VadK in the presence of ATP and Mg_2+_. (B) MS of VadK-H422A alone (black line), with ATP and in the absence (blue line) and presence of Mg_2+_ (red line). No phosphorylation event is observed as there is no mass shift of ∼80 Da in the presence of ATP and Mg_2+_.

Autophosphorylation of canonical PPDK’s occurs at a catalytic histidine in the kinase domain (Herzberg *et al*, 1996) (Minges *et al*, 2017) and we posit this is also the case for VadK. AlphaFold3 (Abramson *et al*., 2024) allowed us to predict the structure of VadK and by overlaying this onto our solved PPDK structures (Fig S2A) we identified a putative autophosphorylation site at His422. To test this prediction, we mutated VadK-His422 to alanine, checked that VadK^His422Ala^ was folded in a similar manner to VadK^WT^ by CD (Fig S4), and repeated the autophosphorylation assay with VadK^WT^ and VadK^His422Ala^. For these experiments, we didn’t remove the His-tag. We showed that the His-tag had no effect on autophosphorylation of VadK^WT^ (Fig S5), however the mutant VadK^His422Ala^ did not autophosphorylate and remained in its apo form (m/z= 52272) (Fig 2B) confirming that His422 is the catalytic residue of VadK. Peptide sequencing of apo- and phosphorylated-VadK by LC-tandem MS analysis confirmed the in silico prediction that His422 was the phosphorylated residue (Fig S6).

### VadK binding sites are essential for growth on propionate

To directly assess the function of the phosphohistidine and ATP binding domains in propionate metabolism we constructed an isogenic panel of *vadK* mutants by complementing Δ*vadK* with a chromosomally integrated copy of *vadK* encoding mutations in the His422 catalytic residue; unphosphorylatable-VadK (Δ*vadK:vadK*^His422Ala^) or phosphomimetic-VadK (Δ*vadK:vadK*^His422Asp^) or the ATP binding site, Δ*vadK:vadK*^Arg106Lys^ (predicted from (Ye *et al*, 2001) to disrupt ATP binding) and compared these strains to WT and Δ*vadK:vadK* (Basu *et al*., 2018).

In standard 7H9 media Δ*vadK*, Δ*vadK:vadK*^His422Ala^, Δ*vadK:vadK*^His422Asp^ and Δ*vadK:vadK*^Arg106Lys^ Mtb strains grew slower than the parental and complemented strain but after 21 days incubation there was no significant differences between the strains (Fig S7A). In 7H9 media with propionate Δ*vadK:vadK*^WT^ grew as WT.

However, the mutant strains expressing unphosphorylatable-Δ*vadK:vadK*^His422Ala^, phosphomimetic Δ*vadK:vadK*^His422Asp^ or the ATP-binding domain mutant Δ*vadK:vadK*^Arg106Lys^ were unable to grow (Fig S7B). These data show that kinase activity is critical for VadK’s role in propionate metabolism as neither the phosphomimetic nor the unphosphorylatable VadK were able to restore VadK growth on propionate.

### VadK does not transcriptionally regulate the MCC during propionate metabolism

Bacterial histidine kinases typically act as two-component regulators that modulate gene expression. To test whether VadK regulates the transcription of genes associated with the MCC we performed global transcriptional profiling. WT and Δ*vad*K were grown to mid-log phase in standard 7H9 media before being washed and inoculated into 7H9 media with propionate. After 4 h incubation, total RNA was isolated and sequenced. These data revealed no significant differences between expression of genes encoding enzymes of the MCC (prpC, prpD, icl1) and MCC transcriptional regulator (prpR) or any other genes associated with CCM in Δ*vadK*, indicating that VadK is not involved in the transcriptional regulation of the MCC or CCM under the conditions tested (DatasetS1). This work indicated that VadK is a non-canonical bacterial histidine kinase.

The RNA-seq analysis did however identify 34 upregulated genes, and 31 downregulated genes (genes differentially expressed log2-fold change ≥±1.2) in Δ*vadK* compared to WT (q<0.001) (DatsetS1). This data was significantly correlated with expression data obtained for Mtb growing in cholesterol (Pawełczyk *et al*., 2021) (chi squared = 1.36^-10^). Of the 34 upregulated genes in Δ*vadK* 15 of these were “cholesterol upregulated” genes (Fig S8). This included *rv1216c*-*rv1218c* operon that encodes an ATP-dependent efflux pump and its transcriptional repressor *raaS* (*rv1219c*) involved in the efflux of acyl-lipids and antibiotics and critical for Mtb long-term survival both in vitro and in vivo (Turapov *et al*, 2014). Δ*vadK* also upregulated the efflux pump *mmpS5*-*mmpL5* and associated transcriptional regulator *mmpR5* (*rv0678*). MmpL5 transporters form functionally redundant complexes with MmpS4 and MmpS5 that together are essential for siderophore export and virulence because in their absence siderophores accumulate intracellularly and poison Mtb (Zhang *et al*, 2020). This operon is also frequently co-expressed with *rv1216c-rv1218c* (Aguilar-Ayala *et al*, 2017).

The operon *rv3269-Rv3270* (*rv3271* is statistically significant but <1.2-fold) is also upregulated. Rv3270 (CtpC) and Rv3269 form a metal efflux system essential for resistance to zinc poisoning and can also efflux multiple antibiotics (Boudehen *et al*, 2022). The other “cholesterol induced” operon is *rv3159c-rv3162c* that similarly is associated with resistance to antibiotics (Aguilar-Ayala *et al*., 2017; Pawełczyk *et al*., 2021).

Of the 31 down regulated genes there were 6 genes that were also down regulated in cholesterol growth conditions. These included *pe15* and *ppe20* that encode a complex that has been shown to transport calcium (Boradia *et al*, 2022). Additionally, there were 7 genes associated with lipid anabolism (*fadA16, fadD9, fadE35, fadE8, fadE5, achA4, rv0315*) that were induced in comparison with the WT parental strain.

### Methylcitrate synthase and methylcitrate dehydratase are interacting partners of VadK

Having established that VadK is not directly controlling the gene expression of the MCC, we then tested the hypothesis that VadK interacts directly with MCC-associated proteins. We performed co-immunoprecipitation experiments using WT Mtb transformed with a plasmid encoding a FLAG-tagged VadK under a constitutive promoter (VadKMtb). We immunoprecipitated from whole cell lysates of WT and VadKMtb grown in standard 7H9 media with and without propionate to identify interactions that are important during growth in propionate. By applying a stringent cut-off to filter out nonspecific binding (proteins with peptide spectrum matches (PSMs) of each replicate ≤1 in the WT control and ≥50 in VadKMtb) we identified interactions between VadK and the MCC enzymes, MCS and MCD (Table 1; DatasetS2). The other top-ranked proteins were all ATP-binding proteins and may represent non-specific associations through conserved nucleotide-binding interfaces rather than bona fide interactions.

**Table 1.** VadK interacts with proteins involved in the methyl citrate cycle.

|  |  |  |  |  |  |
| --- | --- | --- | --- | --- | --- |
| Rank | Rv | Gene | Description | iP:VadkMtb in 7H9 | iP:VadKMtb in 7H9+prop |
| 1 | Rv1131 | *prpC* | 2-methylcitrate synthase | 7.5 | 56 |
| 2 | Rv1130 | *prpD* | 2-methylcitrate dehydratase | 10.5 | 71.5 |
| 3 | Rv1020 | *mfd* | Transcription-repair-coupling factor | 38 | 52 |
| 4 | Rv1179c | *rv1179c* | Helicase ATP-binding domain-containing protein | 46 | 52 |
| 5 | Rv2935 | *ppsE* | Phenolphthiocerol/phthi ocerol polyketide synthase subunit | 78 | 85.5 |
| Mtb transformed with a plasmid encoding VadK-FLAG under a constitutive promoter were grown in standard 7H9 media (7H9) and standard 7H9 media and propionate (7H9 + prop). VadKMtb was immunoprecipitated from whole-cell lysates and interacting proteins were identified by MS. Data were calculated from two biological replicates. Nonspecific binding peptides were removed by setting the filter of Peptide spectrum matches (PSMs) of each replicate to “$\leq 1$” in the WT control samples and “$\geq 50$” in “VadKMtb” samples. Hits were ranked in descending order based on ratio of average PSM’s from VadKMtb grown in 7H9 + Prop versus 7H9. Unfiltered raw counts are available in the Supporting Information (DatasetS2). | | | | | |

### Loss of VadK dysregulates central carbon metabolism, energetics and redox homeostasis in cholesterol/propionate conditions

We previously demonstrated that Δ*vadK* is unable to grow in either cholesterol or propionate even with the addition of an alternative carbon source (Basu *et al*., 2018). To test whether this phenotype is caused by defects in the MCC we applied ^13^C-isotopomer analysis to measure changes in metabolic fluxes using different ^13^C-labelled carbon sources. As *ΔvadK* cannot grow in media containing propionate/cholesterol we grew the strains in growth permissive 7H9 media before washing and then shifting into non-permissive media for 48 h.

Firstly, using [3, 4-^13^C_2_] cholesterol as a tracer and sole carbon source in Roisin’s minimal media we metabolically profiled WT, Δ*vadK* and complement after 48 h (Fig 3). Labelling of intracellular metabolites were measured by MS as previously described (Thomson *et al*., 2022). Cholesterol catabolism yields four propionyl-CoAs, four acetyl-CoAs, one succinyl-CoA and a pyruvate that contains the ^13^C labelling.

**Figure 3.**
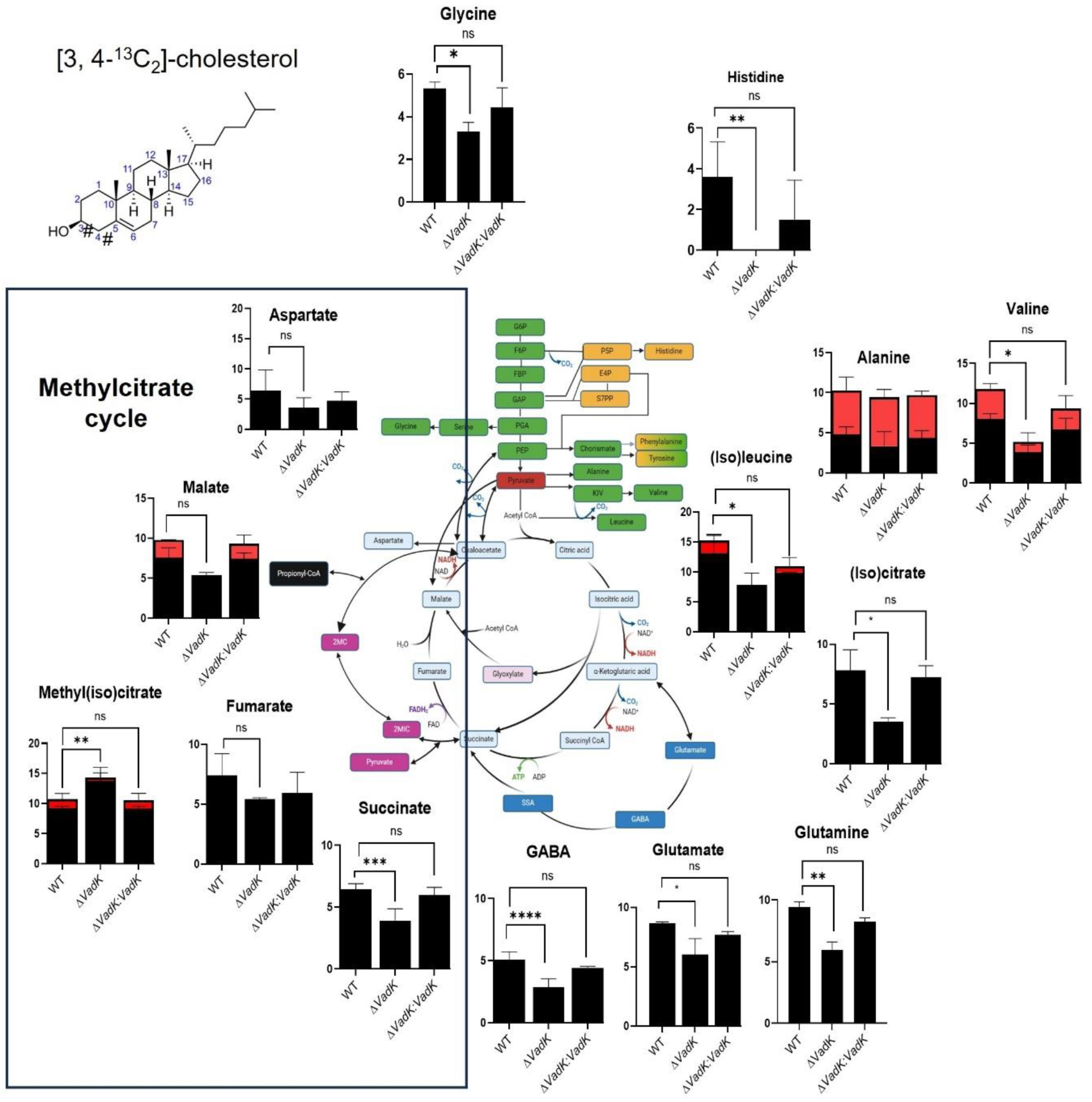
VadK deficient Mtb increases flux through the methylcitrate cycle in cholesterol. ^13^C-isotopomer labelling of metabolites extracted from WT, Δ*vadK* and Δ*vadK:vadK* Mtb strains incubated in Roisin’s media containing 100% [3, 4-^13^C_2_]- labelled cholesterol for 48 h. Average ^13^C-incorporation into 1-carbon (black) and 2-carbons (red) are shown superimposed on a metabolic map. The data represents the mean and SD of three independent biological replicates. P>0.01: NS, P<0.05*, P<0.005**, p<0.0005***, two-way ANOVA with Tukey’s multiple comparison test. # represents the position of the ^13^C label in cholesterol.

The incorporation of [^13^C_2_] into pyruvate derived amino acids alanine and valine accordingly reflects the entry point of the [3, 4-^13^C_2_] cholesterol into metabolism (Fig 3). MCC was active and complete in all three strains as evidenced by ^13^C-incorporation into 2-methyl(iso)citrate (MS cannot distinguish between 2-methylcitrate and 2-methylisocitrate) (Fig 3).

Surprisingly there was significantly increased incorporation of ^13^C into 2-methyl(iso)citrate in Δ*vad*K compared to WT and complemented strain (Fig 3) consistent with enhanced flux through MCS and/or MCD in Δ*vadK* that could potentially generate elevated and toxic concentrations of the MCC intermediates. However, 2-methyl(iso)citrate levels were not significantly different between the strains at this time point (Fig S9).

An alternative explanation is that an overactive MCC consumes OAA and other intermediates required both to fuel the rest of CCM and to grow. In support of this hypothesis there was significantly reduced ^13^C-incorporation into the intermediates of the oxidative branch of the tricarboxylic acid cycle (TCA) cycle ((iso)leucine, (iso)citrate and succinate) in Δ*vadK* and reduced flux through the gamma-aminobutyric acid (GABA) shunt (<^13^C incorporation into glutamate/glutamine and GABA), gluconeogenesis (<^13^C label in glycine) and no flux into the pentose phosphate pathway (PPP) (no ^13^C labelled histidine was detected) as compared with the WT and complement (Fig 3). Overall, these results phenocopy the metabolic profile that we previously reported for intracellular Δ*vadK* within macrophages, conditions where this strain is also unable to grow (Basu *et al*., 2018).

To directly explore the role of VadK in the metabolism of propionate, we performed a mirrored ^13^C labelling experiment. We profiled WT, Δ*vad*K and complement after 48 h of labelling in 7H9 media containing either fully labelled [^13^C_3_]glycerol and unlabelled propionate OR [^13^C_3_]propionate) and unlabelled glycerol. This approach allowed us to calculate how much of the carbon in each metabolite was derived from propionate and how much was derived from glycerol. These experiments demonstrated there were significant differences between the flux of carbon from propionate and glycerol in Δ*vadK* as compared with WT and the complemented strains (Fig 4).

**Figure 4.**
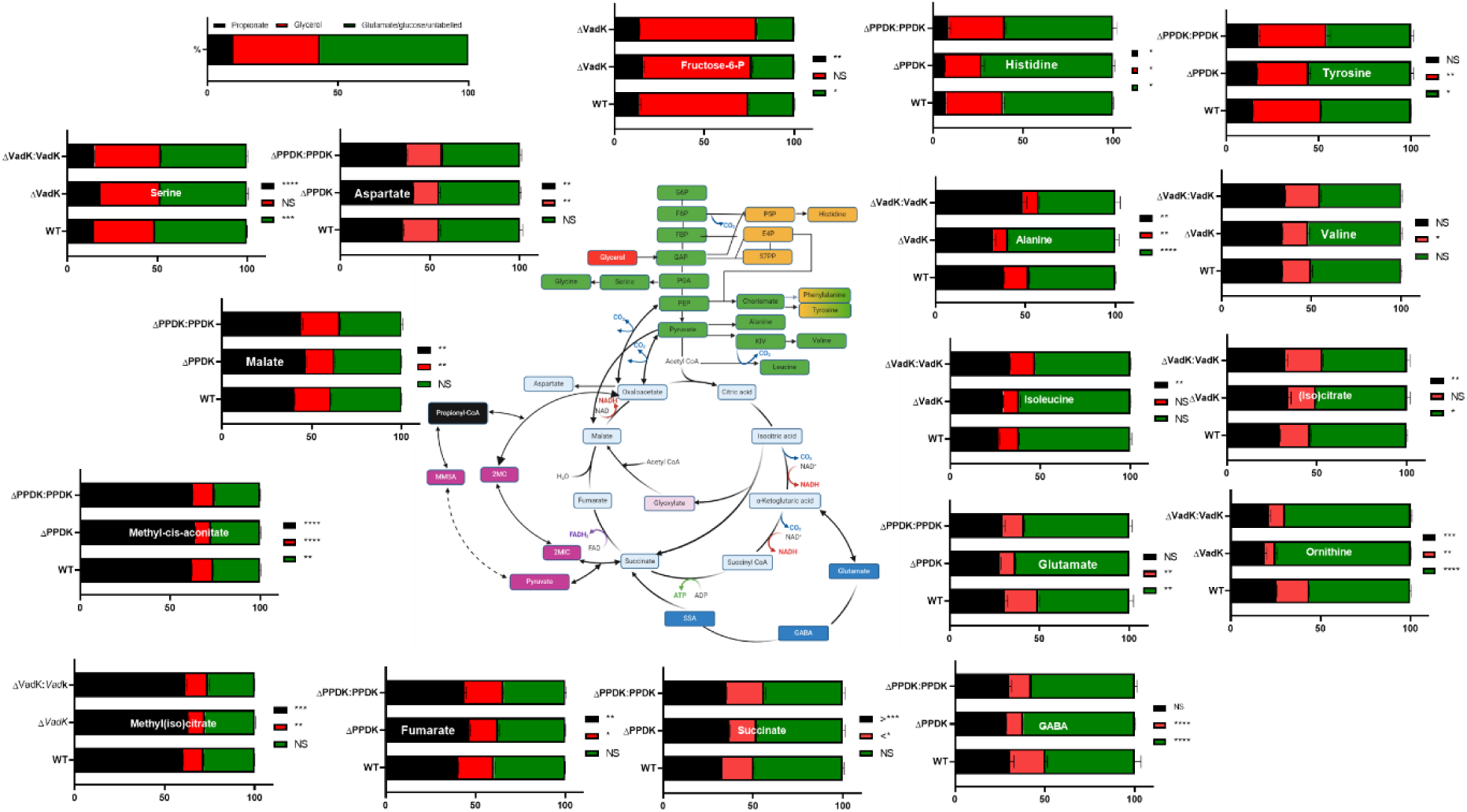
VadK deficient Mtb increases flux of carbon derived from propionate through the methylcitrate cycle. WT, Δ*vadK* and Δ*vadK:vadK* Mtb strains were exposed to 7H9 media (contains unlabelled glutamate and glucose) with either [^13^C_3_]glycerol and unlabelled propionate OR [^13^C_3_]propionate and unlabelled glycerol. The metabolites were extracted and ^13^C labelling measured by mass spectrometry. The percent ^13^C incorporation was used to calculate how much of the carbon from each metabolite was derived from propionate and how much was derived from glycerol. The data represent four experiments (duplicate biological replicates of the two different 13C labelling conditions, total of 4 biological replicates). P>0.05:NS, P<0.05:*, P<0.005, **, p<0.0005:***, p<00005**** two-way ANOVA with Tukey’s multiple comparison test.

All three strains had an active and complete MCC with more than 60% of the carbon in 2-methy(iso)citrate and methyl-cis-aconitate derived from propionate (Fig 4). This shows that even in WT Mtb flux through MCS/MCD is higher than through the rest of the MCC where 32-47% of the carbon backbone from succinate, fumarate, malate and aspartate (as a proxy for oxaloacetate) were derived from propionate (Fig 4). Consistent with the ^13^C profile measured during cholesterol metabolism, loss of vadK led to enhanced flux of propionate through the MCC.

Again, we did not detect differences in 2-methyl(iso)citrate lyase levels suggesting that toxicity doesn’t explain the growth defect. There was notably ∼10-fold more 2-methyl(iso)citrate in propionate versus cholesterol conditions in all Mtb strains tested that could not be explained by differences in amounts of carbon provided (Fig S9). Increased Δv*adK* MCC flux was again correlated with reduced flux into the PPP (Fig 4) but the increased flux to fructose-6-phosphate demonstrated that this was not caused by MCC intermediate–mediated inhibition of fructose-1,6-bisphosphatase as previously described (Eoh & Rhee, 2014).

The flux of carbon from glycerol was significantly reduced in Δ*vadK* in comparison to WT and complement except to fructose 6-phosphate, serine and (iso)citrate where no significant change was measured (Fig 4). This suggested that increased flux of propionate through the MCC in Δ*vadK* was affecting glycerol metabolism more generally in the mutant.

Kinetic modelling predicts that MCS controls the catabolic flux though the MCC (Tummler *et al*, 2018). To validate the increase in flux through the MCC, given that MCS was identified as an interacting partner of VadK, we measured MCS activity in cell-free extracts of WT, Δ*vadK* and Δ*vadK:vadK*. To do this we performed carbon shift experiments as described (Muñoz-Elías *et al*., 2006) in which Mtb strains were grown to mid-log phase in standard 7H9 media, then shifted into 7H9 media plus propionate and incubated for 1 or 4 days before MCS activities were measured (Fig 5). These data showed that in accordance with the flux data Δ*vadK* had significantly increased MCS activity after 1 (303%) and 4 days (447%) as compared with the wildtype and complemented strain. These data confirm the increased flux through the MCC and provide further evidence that VadK is regulating the MCC.

**Figure 5.**
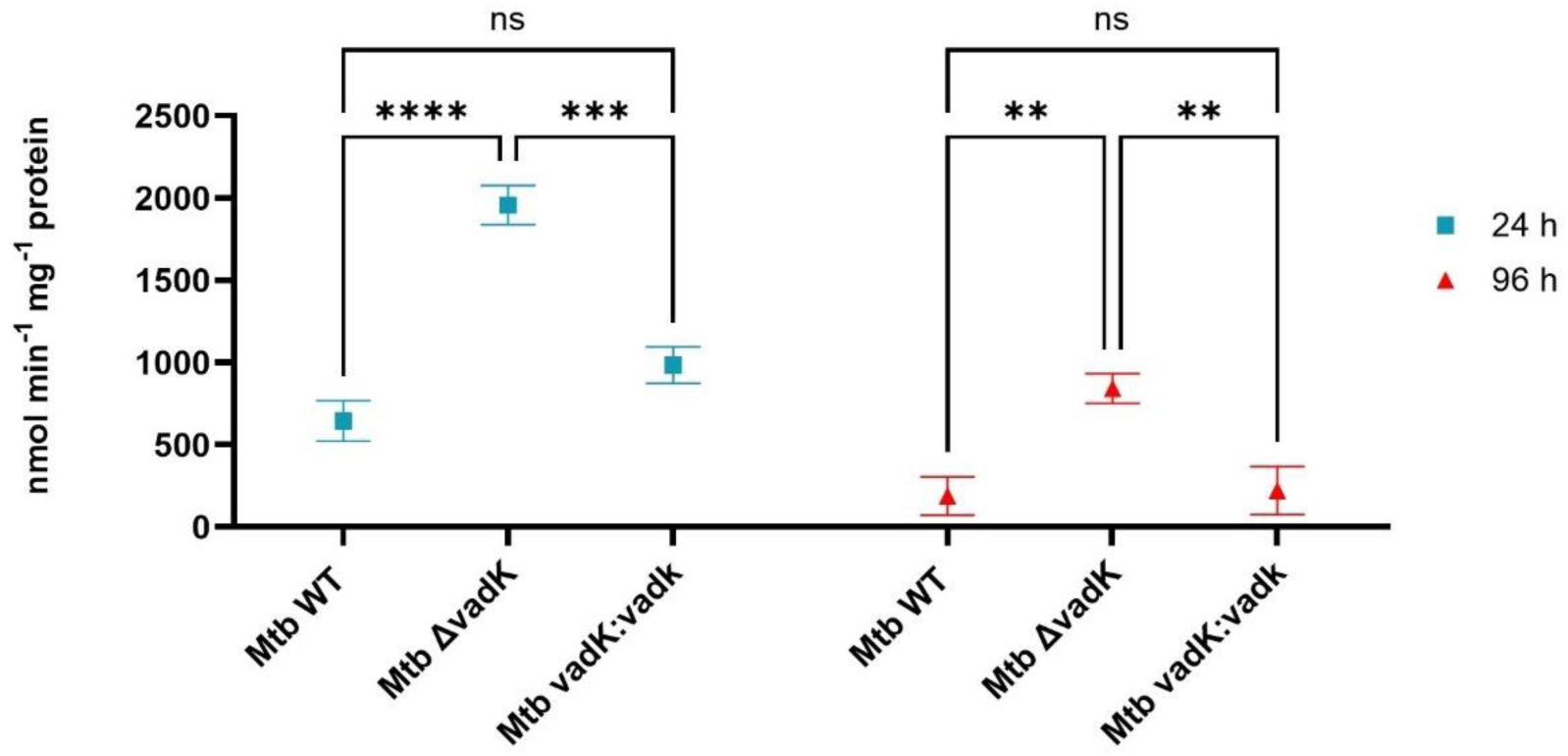
VadK deficient Mtb has significantly enhanced methylcitrate synthase activity. WT, Δ*vadK*, Δ*vadK:vadk* were grown to mid-log in 7H9 media then shifted into 7H9 media containing 10 mM propionate for 1 or 4 days before MCS activity was measured. The average ±SEM rate of CoA production from 3 independent biological replicates is shown. P>0.05:NS, P<0.05:*, P<0.005, **, p<0.0005:***, p<00005**** two-way ANOVA with Tukey’s multiple comparison test.

To complement the metabolic flux analysis, Seahorse extracellular flux assays were performed to measure cellular bioenergetics of WT, Δ*vadK* and Δ*vadK:vadK* in 7H9 media with propionate. The bioenergetic phenogram indicated that ΔvadK has an energetically quiescent phenotype in concordance with reduced TCA flux and no growth in these conditions (Fig S10A-C).

Given that reduced flux through both the oxidative branch of the TCA cycle and the pentose phosphate pathway in Δ*vadK*, we hypothesized that cellular redox homeostasis could be perturbed in the mutant. To test this hypothesis, we used a previously described redox biosensor (Mrx1-roGFP2) that measures the redox potential of the antioxidant buffer mycothiol (reduced [MSH]/oxidized [MSSM]; EMSH) as a quantifiable indicator of cytoplasmic redox state of Mtb (Bhaskar *et al*, 2014). An increase in the 405/488 ratio indicates an oxidative shift in the MSH/MSSM ratio. As shown (Fig S10D) the intramycobacterial redox state of Δv*adK* was significantly more oxidized than either the WT or complemented strain of Mtb in propionate conditions which is concordance with ^13^C data showing the reduction of flux through the PPP and oxidative arm of the TCA cycle (Fig 4).

### ATP accumulates in Δ*vadK*

Given that Δ*vadK* perturbs CCM and that VadK requires ATP for activity, we measured ATP levels in Mtb strains grown in 7H9 medium with or without propionate (Fig S11). No significant differences in ATP were observed among WT, Δ*vadK*, and complemented strains in standard 7H9 medium. In propionate-containing media, ATP levels were significantly elevated in all strains after 7 days (Fig S11). This was not due to an artifact of the extraction process (Mulholland *et al*, 2025) as the cells were lysed by bead beating (methods). By 14 days, WT and Δ*vadK:vadK* strains returned to levels comparable to propionate-free conditions. In contrast, Δ*vadK* maintained higher ATP levels than the WT and complement strains.

### Loss of VadK does not affect cell wall lipid biosynthesis

As propionyl-CoA can also be channelled directly into the biosynthesis of cell wall lipids we extracted lipids from WT, Δ*vadK* and Δ*vadK:vadK* grown in 7H9 media with propionate and analysed them by 2D-TLC to test the effects of VadK deletion on cell wall lipids (Fig S12A). There were no significant differences in PDIM incorporation between WT, Δ*vadK* and Δ*vadK:vadK* which is accordance with previous studies that showed that incorporation of carbon from propionate into PDIM’s is a constitutive process, occurring independently of the amount of flux through the MCC (Lee *et al*., 2013). Production of the sulfolipid SL-1, was however slightly elevated in the mutant strain as compared to the WT and complemented strains (Fig S12B).

### VadK-deficient Mtb is attenuated for growth on glycerol and pyruvate

The role of the canonical MCC pathway is well established for the catabolism of propionate and cholesterol in Mtb and other bacteria and fungi (Dolan *et al*, 2018) (Huang *et al*, 2023). However, Serafini *et al* (2019) (Serafini *et al*, 2019) demonstrated that Mtb can operate a reversed MCC when growing on pyruvate or lactate. We similarly showed that this was also the case for glycerol-grown Mtb (Borah *et al*., 2021). Here, ^13^C-metabolic flux analysis also demonstrated that Δ*vad*K had dysregulated glycerol metabolism (Fig 4). To test the hypothesis that VadK is important for growth on carbon sources that require a reversed MCC we cultured WT, Δ*vadK* and Δ*vad*K:*vad*K in Roisin’s minimal media containing either glycerol or pyruvate as sole carbon sources (Fig 6). These data showed that Δ*vad*K was significantly attenuated for growth on either of these carbon sources as compared with the WT and complement (Fig 6). To confirm that this phenotype is a consequence of a dysregulated MCC we tested whether B_12_ chemically complemented the growth phenotype. Vitamin B_12_ downregulates the expression of *prpR*-*prpD*-*prpC* operon in Mtb effectively turning the MCC off (Campos-Pardos *et al*, 2024) and activates the methylmalonyl pathway because it’s an essential cofactor for the second enzyme in this pathway, methylmalonyl-CoA mutase (MCM). In accordance with our hypothesis vitamin B_12_ partially rescued the growth phenotype of Δ*vadK* in these carbon sources (Fig 6).

**Figure 6.**
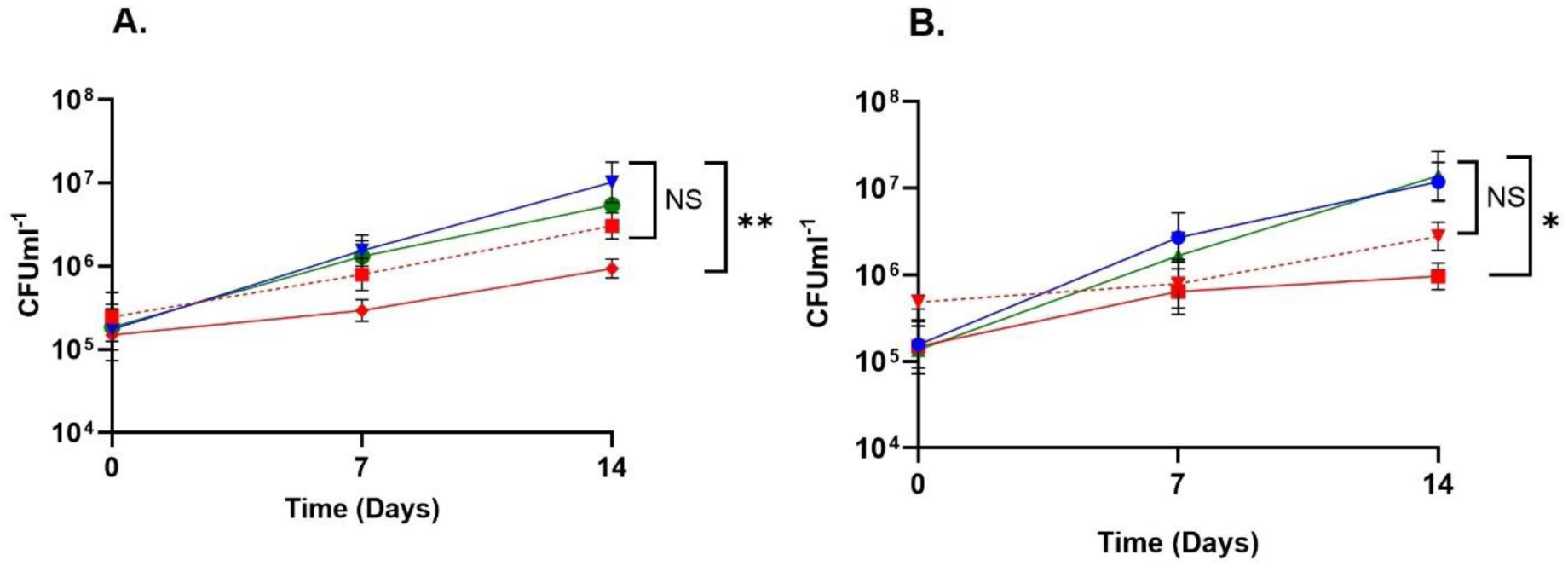
VadK deficient Mtb is attenuated for growth on pyruvate and glycerol. WT (blue) and Δ*vadK* (red), Δ*vadK:vadk* (green) Mtb strains were grown in Roisin’s minimal media containing either (A) glycerol or (B) pyruvate as sole carbon source. ΔvadK is partially rescued by the addition of vitamin B_12_ (red dotted line). Growth was measured by enumerating the CFU’s and the results represent the mean and SEM of x 2-3 biological replicates. P>0.05:NS, P<0.05:*, two-way ANOVA with Tukey’s multiple comparison test.

### PPDK has evolved into VadK in all pathogenic mycobacteria and related genera

Having established the significant role of VadK in the metabolism and virulence of Mtb we wanted to explore the distribution of VadK across mycobacteria and related genera. This analysis (Fig 7) showed that in addition to Mtb VadK is present in pathogenic mycobacteria including the causative agent of leprosy (*Mycobacterium leprae*), bovine TB (*Mycobacterium bovis*), Buruli Ulcer (*Mycobacterium ulcerans*) and within *Mycobacterium abscesses*-an important pathogen in people with chronic lung disease. However, non-pathogens such as *Mycobacterium smegmatis* completely lacks VadK whilst other mycobacteria have full length PPDK but lack critical motifs for kinase activity (*Mycobacterium paraseoulense*, and *Mycobacterium kiuyosense*). Additionally, VadK was identified in the genome of other actinomyces including Nocardia, Streptomyces and Rhodococcus spp. indicating that vadK has evolved multiple times from PPDK or has been acquired by horizontal gene transfer.

**Figure 7.**
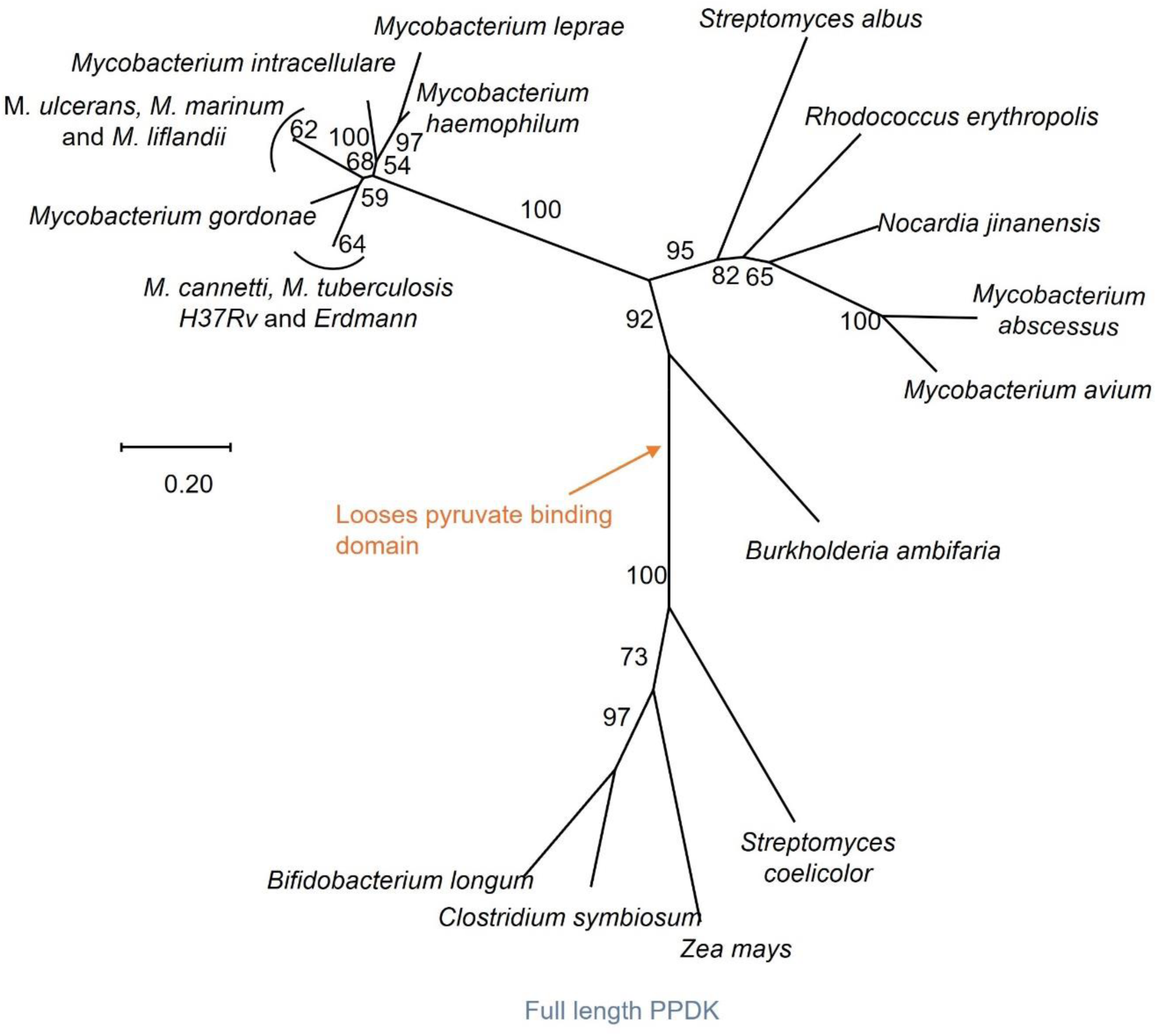
VadK has evolved independently multiple times. The evolutionary history was inferred with MEGAX(Kumar *et al*., 2018) using (Whelan & Goldman, 2001). The percentage of trees in which the associated taxa clustered together is shown next to the branches. Branch lengths indicate the number of substitutions per site. All positions containing gaps and missing data were eliminated.

## Discussion

Phosphorylation is a widespread mechanism used to regulate protein function during adaptation to different physiological conditions in the host. In bacteria this is typically mediated via protein kinases that phosphorylate serine, tyrosine or threonine or histidine/aspartate based two component regulators. We have discovered a unique, non-canonical histidine (di)kinase, VadK that has evolved from an ancestral PPDK in pathogenic mycobacteria including Mtb and other related actinomyces. We have demonstrated that VadK is a functional kinase that carries out a regulatory role in the MCC that is essential for Mtb growth and virulence in animal models of tuberculosis.

Metabolism of propionate via the MCC is a metabolic strategy that is critical for the virulence and survival of several pathogens in the host (Dolan *et al*., 2018). Propionate can be derived from the metabolism of host-derived carbon sources such as sterols, branched chain amino acids and fatty acids and is also present at high (mM) levels during certain diseases such as the lungs of cystic fibrosis patients (Ghorbani *et al*, 2015). However, flux through the MCC presents microbes with a challenge. MCC intermediates can inhibit microbial growth, and the pathway consumes intermediates of the TCA cycle that must be replenished through anaplerotic pathways. Therefore, the MCC must be carefully regulated to prevent collateral harm to the rest of CCM.

Some of the first studies to explore the role of the MCC in metabolism and virulence of microbes were performed using strains of Mtb with deletions that interrupted the MCC (Muñoz-Elías *et al*., 2006). These strains were growth attenuated in media containing either propionate or cholesterol (Muñoz-Elías *et al*., 2006). Although the exact mechanism of growth inhibition remains unknown, propionate toxicity is correlated with reduced flux through the TCA, reduced energy production and inhibition of the PPP in Mtb (Dolan *et al*., 2018; Eoh & Rhee, 2014).

Here, we have identified that Δ*vadK* has a growth and metabolic phenotype strikingly similar to that of MCC-deficient strains but without any lesion in the MCC. Mtb strains lacking VadK exhibit increased flux through the MCC that prevents in vitro growth of Mtb in propionate/cholesterol containing media and attenuates growth on media with glycerol or pyruvate as the sole carbon source. These phenotypes are driven by the impact of increased flux through the MCC that competes for TCA-intermediates reducing flux through the oxidative branch of the TCA cycle and through the PPP causing changes in redox balance.

RNA-seq analysis demonstrated that VadK is not transcriptionally regulating the MCC and showed a significant overlap with the transcriptomic response to cholesterol, specifically the upregulation of detoxification systems and efflux of lipids/antibiotics/mycobactin and the down regulation of genes involved in amino acid and lipid metabolism as well as calcium import. These transcriptional signatures correlate to increased ATP under these conditions as calcium and ATP levels are intrinsically linked. Calcium transport is dependent on ATP, but its import also enhances ATP production.

Interestingly this work demonstrated that levels of 2-methyl(iso)citrate are significantly higher when Mtb is exposed directly to propionate versus substrates that yield propionyl CoA such as cholesterol. The regulatory mechanisms underpinning these differences require further investigations. This is significant as it has been shown that Mtb has a mixed diet in the host which includes propionyl CoA yielding carbon sources such as sterols whereas *M. abscessus* within a cystic fibrosis lung has direct access to propionyl CoA (Beste *et al*, 2013) (Dolan *et al*., 2018).

We previously demonstrated that VadK is required for the growth of Mtb in macrophages (Basu *et al*., 2018) and here we extended this work to show that it is also required for virulence in two different murine models of tuberculosis. The connection between the MCC and virulence has been observed previously. For example, blocking the MCC impairs the virulence of several pathogenic fungi and bacteria (Dolan *et al*., 2018). The MCC is necessary for virulence of the rice blast fungus, *Pyricularia oryzae* (Yan *et al*, 2019) and *Aspergillus fumigatus* (Ibrahim-

Granet *et al*, 2008) and is important to *Neisseria meningitidis* infection and transmission (Catenazzi *et al*, 2014). By contrast whilst deleting MCL activity (encoded by *icl* genes) or *prpDC* from Mtb impaired growth in macrophages, only the MCL mutant was attenuated *in vivo* (Muñoz-Elías & McKinney, 2005). Interpretation of this result is complicated in Mtb as the MCL’s also function as isocitrate lyase’s in the glyoxylate shunt, so deleting the MCL ablates both pathways. Elegant studies by Rhee *et al* (2014) indicated that *in vitro* the defects associated with icl deficiency are due to a broken MCC. Breaking the MCC at MCL leads to the build-up of the toxic intermediate 2-methylisocitrate (Eoh & Rhee, 2014). It is presumed that in vivo, *prpDC* is dispensable because there is sufficient B_12_ to activate the B_12_-dependent methyl malonyl pathway to metabolise propionate. Vitamin B_12_ can also down regulate the MCC by repressing the transcription of *prpCD* and the MCC transcriptional regulator *prpR* (Campos-Pardos *et al*., 2024). In contrast to disrupting the MCC, we show that dysregulating flux through the MCC severely attenuates Mtb in vivo and so represents a better strategy to target Mtb.

The requirement for VadK for virulence and survival in the murine models suggests that in vivo B_12_ concentrations do not fully repress the MCC and/or that VadK regulates MCC, but via a mechanism other than transcription. Although murine serum B_12_ levels vary significantly they are generally in the picogram range rather than the microgram levels of B_12_ required to rescue Δ*vadK* (Campos-Pardos *et al*., 2024). In vivo data indicates that while B_12_ concentrations are adequate to induce the methylmalonyl pathway, they are not high enough to completely suppress the MCC (Campos-Pardos *et al*., 2024). As an extension of our work, it would be interesting to test the phenotype of Δ*vadK* in a B_12_ anaemic murine model of tuberculosis. Future studies will be required to establish the reach of VadK’s regulatory effects.

It was previously established that a reverse MCC pathway is utilised by Mtb grown on pyruvate, lactate (Serafini *et al*., 2019) and glycerol (Borah *et al*., 2021). However, *prpDC* is not essential for growth in either pyruvate or lactate demonstrating that although the MCC is active under these conditions it is not essential (Serafini *et al*., 2019). By contrast, we have demonstrated that deletion of VadK attenuates the growth of Mtb in either glycerol or pyruvate and that this phenotype is rescued by supplementation with B_12_, demonstrating that downregulation of the MCC and/or activation of the methyl malonyl pathway restores growth of this mutant.

Overall, this work demonstrates that dysregulation of the MCC caused by the loss of VadK has broad impacts on Mtb metabolism and growth than complete loss of the pathway, and links MCC hyperactivity to a slow- or non-replicating phenotype. These findings raise the intriguing hypothesis that fine tuning flux through the MCC could serve as another mechanism by which Mtb regulates its growth rate. Given that VadK’s autophosphorylation requires ATP, it could function as a sensor of cellular energy state: high ATP levels driving VadK-mediated suppression of MCC flux, while low ATP could enhance MCC flux, imposing a brake/slowdown of growth. Supporting this hypothesis, we demonstrated that ATP levels increase in response to propionate even in WT Mtb but this is reduced over time. In the absence of VadK, however ATP levels remained elevated. We and others have previously demonstrated that ATP is growth inhibitory to Mycobacteria, and this therefore could be contributing to the growth attenuation of Δ*vadK* (Lodhiya *et al*, 2024; Tatano *et al*, 2015). Further work is required to test this hypothesis.

Our phylogenetic trees indicate that VadK orthologues are conserved across pathogenic mycobacteria -including amongst *M. leprae’s* minimal set of proteins, supporting a key role for VadK in mycobacterial pathogenesis and its potential as a drug target. As propionate accumulates in the lungs during certain diseases such as the lungs of cystic fibrosis patients at millimolar concentrations (Ghorbani *et al*., 2015) careful regulation of flux through the MCC is likely to be important to various pathogens that cause disease in this environment. Notably, non-tuberculous mycobacteria, including *M. abscessus* that causes the most severe disease and highest mortality in individuals with chronic lung conditions such as chronic obstructive pulmonary disease, cystic fibrosis, and bronchiectasis encodes a VadK orthologue. We also identified VadK in *Burkholderia ambifaria*, a plant associated bacterium that causes opportunistic infections in humans, particularly in cystic fibrosis patients (Vinacour *et al*, 2023), suggesting VadK may contribute more broadly to bacterial virulence, particularly in hosts with chronic lung disease.

In summary, we have identified a novel kinase, VadK, that is essential to cause tuberculosis and for Mtb growth on carbon sources metabolized through the MCC and to cause tuberculosis. VadK likely functions to fine-tune flux through the methylcitrate cycle, so coordinating MCC activity with the rest of the CCM and the cells energy state. The presence of VadK orthologues in other mycobacterial and non-mycobacterial pathogens suggests that this kinase plays a role in the pathogenesis of organisms that rely on the MCC within the host. Importantly, VadK has no homologues in humans, highlighting its potential as a candidate for future drug development.

## Acknowledgements

Thanks to the funders of this work: Biotechnology and Biological Sciences Research Council (BB/T007648/1; BB/V018159/1), the Medical Research Council (MR/W018756/1), National Institutes of Health (T32 AI141346; AI095208), Department of Biotechnology (BT/PR47905/MED/29/ 1643/2023).

Special thanks to Kate Heesom for all the helpful proteomic analysis. We thank Dirk Schnappinger and Sabine Ehrt (Weill Cornell Medical College, New York) for kindly providing the cloning vectors used in the Gateway assembly of the Rv1127c expression plasmid.

## References

Abramson J, Adler J, Dunger J, Evans R, Green T, Pritzel A, Ronneberger O, Willmore L, Ballard AJ, Bambrick J et al (2024) Accurate structure prediction of biomolecular interactions with AlphaFold 3. Nature 630: 493–500

Aguilar-Ayala DA, Tilleman L, Van Nieuwerburgh F, Deforce D, Palomino JC, Vandamme P, Gonzalez YMJA, Martin A (2017) The transcriptome of Mycobacterium tuberculosis in a lipid-rich dormancy model through RNAseq analysis. Sci Rep 7: 17665

Basu P, Sandhu N, Bhatt A, Singh A, Balhana R, Gobe I, Crowhurst NA, Mendum TA, Gao L, Ward JL et al (2018) The anaplerotic node is essential for the intracellular survival of *Mycobacterium tuberculosis*. J Biol Chem 293: 5695–5704

Beste DJ, Laing E, Bonde B, Avignone-Rossa C, Bushell ME, McFadden JJ (2007) Transcriptomic analysis identifies growth rate modulation as a component of the adaptation of mycobacteria to survival inside the macrophage. J Bacteriol 189: 3969–3976

Beste DJ, Noh K, Niedenfuhr S, Mendum TA, Hawkins ND, Ward JL, Beale MH, Wiechert W, McFadden J (2013) 13C-flux spectral analysis of host-pathogen metabolism reveals a mixed diet for intracellular *Mycobacterium tuberculosis*. Chem Biol 20: 1012–1021

Beste DJ, Peters J, Hooper T, Avignone-Rossa C, Bushell ME, McFadden J (2005) Compiling a molecular inventory for *Mycobacterium bovis* BCG at two growth rates: evidence for growth rate-mediated regulation of ribosome biosynthesis and lipid metabolism. J Bacteriol 187: 1677–1684

Beste DJV, Espasa M, Bonde B, Kierzek AM, Stewart GR, McFadden J (2009) The genetic requirements for fast and slow growth in mycobacteria. PloS one 4: e5349–e5349

Bhaskar A, Chawla M, Mehta M, Parikh P, Chandra P, Bhave D, Kumar D, Carroll KS, Singh A (2014) Reengineering Redox Sensitive GFP to Measure Mycothiol Redox Potential of *Mycobacterium tuberculosis* during Infection. PLOS Pathog 10: e1003902

Boradia V, Frando A, Grundner C (2022) The *Mycobacterium tuberculosis* PE15/PPE20 complex transports calcium across the outer membrane. PLoS Biol 20: e3001906

Borah K, Mendum TA, Hawkins ND, Ward JL, Beale MH, Larrouy-Maumus G, Bhatt A, Moulin M, Haertlein M, Strohmeier G et al (2021) Metabolic fluxes for nutritional flexibility of *Mycobacterium tuberculosis*. Mol Syst Biol 17: e10280

Boudehen YM, Faucher M, Maréchal X, Miras R, Rech J, Rombouts Y, Sénèque O, Wallat M, Demange P, Bouet JY et al (2022) Mycobacterial resistance to zinc poisoning requires assembly of P-ATPase-containing membrane metal efflux platforms. Nat Commun 13: 4731

Burley KH, Cuthbert BJ, Basu P, Newcombe J, Irimpan EM, Quechol R, Foik IP, Mobley DL, Beste DJV, Goulding CW (2021) Structural and Molecular Dynamics of *Mycobacterium tuberculosis* Malic Enzyme, a Potential Anti-TB Drug Target. ACS Infect Dis 7: 174–188

Campos-Pardos E, Uranga S, Picó A, Gómez AB, Gonzalo-Asensio J (2024) Dependency on host vitamin B12 has shaped *Mycobacterium tuberculosis* Complex evolution. Nat Commun 15: 2161

Catenazzi MC, Jones H, Wallace I, Clifton J, Chong JP, Jackson MA, Macdonald S, Edwards J, Moir JW (2014) A large genomic island allows *Neisseria meningitidis* to utilize propionic acid, with implications for colonization of the human nasopharynx. Mol Microbiol 93: 346–355

Consortium TU (2024) UniProt: the Universal Protein Knowledgebase in 2025. Nucl Acids Res 53: D609–D617

Das M, Sreedharan S, Shee S, Malhotra N, Nandy M, Banerjee U, Kohli S, Rajmani RS, Chandra N, Seshasayee ASN et al (2023) Cysteine desulfurase (IscS)-mediated fine-tuning of bioenergetics and SUF expression prevents *Mycobacterium tuberculosis* hypervirulence. Sci Adv 9: eadh2858

de Miranda R, Cuthbert BJ, Klevorn T, Chao A, Mendoza J, Arbing M, Sieminski PJ, Papavinasasundaram K, Abdul-Hafiz S, Chan S et al (2023) Differentiating the roles of *Mycobacterium tuberculosis* substrate binding proteins, FecB and FecB2, in iron uptake. PLoS Pathog 19: e1011650

Dobson G, Minnikin D, Goodfellow M (1983). Systematic analysis of complex mycobacterial lipids pp. R18–R18.

Dolan SK, Wijaya A, Geddis SM, Spring DR, Silva-Rocha R, Welch M (2018) Loving the poison: the methylcitrate cycle and bacterial pathogenesis. Microbiol 164: 251–259

Ehrt S, Guo XV, Hickey CM, Ryou M, Monteleone M, Riley LW, Schnappinger D (2005) Controlling gene expression in mycobacteria with anhydrotetracycline and Tet repressor. Nucleic Acids Res 33: e21

Eoh H, Rhee KY (2014) Methylcitrate cycle defines the bactericidal essentiality of isocitrate lyase for survival of *Mycobacterium tuberculosis* on fatty acids. Proc Natl Acad Sci U S A 111: 4976–4981

Ghorbani P, Santhakumar P, Hu Q, Djiadeu P, Wolever TMS, Palaniyar N, Grasemann H (2015) Short-chain fatty acids affect cystic fibrosis airway inflammation and bacterial growth. Eur Resp J 46: 1033–1045

Griffin JE, Pandey AK, Gilmore SA, Mizrahi V, McKinney JD, Bertozzi CR, Sassetti CM (2012) Cholesterol catabolism by *Mycobacterium tuberculosis* requires transcriptional and metabolic adaptations. Chem Biol 19: 218–227

Herzberg O, Chen CC, Kapadia G, McGuire M, Carroll LJ, Noh SJ, Dunaway-Mariano D (1996) Swiveling-domain mechanism for enzymatic phosphotransfer between remote reaction sites. PNAS 93: 2652–2657

Huang Z, Wang Q, Khan IA, Li Y, Wang J, Wang J, Liu X, Lin F, Lu J (2023) The methylcitrate cycle and its crosstalk with the glyoxylate cycle and tricarboxylic acid cycle in pathogenic fungi. Molecules 28

Ibrahim-Granet O, Dubourdeau M, Latgé JP, Ave P, Huerre M, Brakhage AA, Brock M (2008) Methylcitrate synthase from *Aspergillus fumigatus* is essential for manifestation of invasive aspergillosis. Cell Microbiol 10: 134–148

Kim JH, Wei JR, Wallach JB, Robbins RS, Rubin EJ, Schnappinger D (2011) Protein inactivation in mycobacteria by controlled proteolysis and its application to deplete the beta subunit of RNA polymerase. Nucleic Acids Res 39: 2210–2220

Koendjbiharie JG, van Kranenburg R, Kengen SWM (2020) The PEP-pyruvate-oxaloacetate node: variation at the heart of metabolism. FEMS Microbiol Reviews 45

Kramnik I, Beamer G (2016) Mouse models of human TB pathology: roles in the analysis of necrosis and the development of host-directed therapies. Semin Immunopathol 38: 221–237

Kramnik I, Dietrich WF, Demant P, Bloom BR (2000) Genetic control of resistance to experimental infection with virulent Mycobacterium tuberculosis PNAS 97: 8560–8565

Kumar S, Stecher G, Li M, Knyaz C, Tamura K (2018) MEGA X: Molecular evolutionary genetics analysis across computing platforms. Mol Biol Evol 35: 1547–1549

Lakshmanan M, Xavier AS (2013) Bedaquiline - The first ATP synthase inhibitor against multi drug resistant tuberculosis. J Young Pharm 5: 112–115

Lee W, VanderVen BC, Fahey RJ, Russell DG (2013) Intracellular *Mycobacterium tuberculosis* exploits host-derived fatty acids to limit metabolic stress. J Biol Chem 288: 6788–6800

Lodhiya T, Palande A, Veeram A, Larrouy-Maumus G, Beste DJV, Mukherjee R, 2024. ATP burst is the dominant driver of antibiotic lethality in Mycobacteria. eLife Sciences 13:RP99656.

Mackenzie JS, Lamprecht DA, Asmal R, Adamson JH, Borah K, Beste DJV, Lee BS, Pethe K, Rousseau S, Krieger I et al (2020) Bedaquiline reprograms central metabolism to reveal glycolytic vulnerability in *Mycobacterium tuberculosis*. Nat Comm 11: 6092

Meade RK, Long JE, Jinich A, Rhee KY, Ashbrook DG, Williams RW, Sassetti CM, Smith CM (2023) Genome-wide screen identifies host loci that modulate *Mycobacterium tuberculosis* fitness in immunodivergent mice. G3 (Bethesda). 2023 Aug 30;13(9):jkad147.

Minges A, Ciupka D, Winkler C, Höppner A, Gohlke H, Groth G (2017) Structural intermediates and directionality of the swiveling motion of Pyruvate Phosphate Dikinase. Sci Rep 7: 45389

Mulholland CV, Lee BS, Chong A, Cui J, Pethe K, Berney M (2025) The mycobacterial ATP burst is a lysis artifact and serves as an assay for drug-induced cell wall damage. bioRxiv: 2025.2010.2024.684413

Muñoz-Elías EJ, McKinney JD (2005) *Mycobacterium tuberculosis* isocitrate lyases 1 and 2 are jointly required for in vivo growth and virulence. Nat Med 11: 638–644

Muñoz-Elías EJ, Upton AM, Cherian J, McKinney JD (2006) Role of the methylcitrate cycle in *Mycobacterium tuberculosis* metabolism, intracellular growth, and virulence. Mol Microbiol 60: 1109–1122

Om K, Arias NN, Jambor CC, MacGregor A, Rezachek AN, Haugrud C, Kunz HH, Wang Z, Huang P, Zhang Q et al (2022) Pyruvate, phosphate dikinase regulatory protein impacts light response of C4 photosynthesis in Setaria viridis. Plant Physiol 190: 1117–1133

Pawełczyk J, Brzostek A, Minias A, Płociński P, Rumijowska-Galewicz A, Strapagiel D, Zakrzewska-Czerwińska J, Dziadek J (2021) Cholesterol-dependent transcriptome remodeling reveals new insight into the contribution of cholesterol to *Mycobacterium tuberculosis* pathogenesis. Sci Rep 11: 12396

Russell DG, VanderVen BC, Lee W, Abramovitch RB, Kim MJ, Homolka S, Niemann S, Rohde KH (2010) *Mycobacterium tuberculosis* wears what it eats. Cell Host Microbe 8: 68–76

Serafini A, Tan L, Horswell S, Howell S, Greenwood DJ, Hunt DM, Phan MD, Schembri M, Monteleone M, Montague CR et al (2019) *Mycobacterium tuberculosis r*equires glyoxylate shunt and reverse methylcitrate cycle for lactate and pyruvate metabolism. Mol Microbiol 112: 1284–1307

Smith CM, Baker RE, Proulx MK, Mishra BB, Long JE, Park SW, Lee H-N, Kiritsy MC, Bellerose MM, Olive AJ et al (2022) Host-pathogen genetic interactions underlie tuberculosis susceptibility in genetically diverse mice. eLife 11: e74419

Tatano Y, Kanehiro Y, Sano C, Shimizu T, Tomioka H (2015) ATP Exhibits Antimicrobial Action by Inhibiting Bacterial Utilization of Ferric Ions. Sci Rep 5: 8610

Thomson M, Liu Y, Nunta K, Cheyne A, Fernandes N, Williams R, Garza-Garcia A, Larrouy-Maumus G (2022) Expression of a novel mycobacterial phosphodiesterase successfully lowers cAMP levels resulting in reduced tolerance to cell wall targeting antimicrobials. J Biolog Chem 298

Tummler K, Zimmermann M, Schubert OT, Aebersold R, Kühn C, Sauer U, Klipp E (2018) Two parallel pathways implement robust propionate catabolism and detoxification in mycobacteria. bioRxiv: 258947

Turapov O, Waddell SJ, Burke B, Glenn S, Sarybaeva AA, Tudo G, Labesse G, Young DI, Young M, Andrew PW et al (2014) Oleoyl coenzyme A regulates interaction of transcriptional regulator RaaS (Rv1219c) with DNA in mycobacteria. J Biol Chem 289: 25241–25249

Vinacour M, Moiana M, Forné I, Jung K, Bertea M, Valdayo PMC, Nikel PI, Imhof A, Palumbo MC, Porto DFD et al (2023) Genetic dissection of the degradation pathways for the mycotoxin fusaric acid in *Burkholderia ambifaria* T16. App Environ Microbiol 89: e00630–00623

Whelan S, Goldman N (2001) A General Empirical Model of Protein Evolution Derived from Multiple Protein Families Using a Maximum-Likelihood Approach. Mol Biol Evolution 18: 691–699

World Health Oraganisation, (2024). Global Tuberculosis Report 2024. World Health Organization, Geneva.

Yan Y, Wang H, Zhu S, Wang J, Liu X, Lin F, Lu J (2019) The Methylcitrate Cycle is Required for Development and Virulence in the Rice Blast Fungus *Pyricularia oryzae*. Mol Plant Microbe Interact 32: 1148–1161

Ye D, Wei M, McGuire M, Huang K, Kapadia G, Herzberg O, Martin BM, Dunaway-Mariano D (2001) Investigation of the Catalytic site within the ATP-grasp domain of *Clostridium symbiosum* Pyruvate Phosphate Dikinase*. J Biol Chem 276: 37630–37639

Zhang L, Hendrickson RC, Meikle V, Lefkowitz EJ, Ioerger TR, Niederweis M (2020) Comprehensive analysis of iron utilization by *Mycobacterium tuberculosis*. PLoS Pathog 16: e1008337

