## Supplementary figures for "VadK, a Non-Canonical Kinase that Regulates the Methylcitrate Cycle and is Essential for *Mycobacterium tuberculosis* Virulence"

**A.**

Reaction 1. *Enzyme + ATP + Pi ↔ Enzyme – His – P + AMP + PP*

Reaction 2. *Enzyme – His – P + pyruvate ↔ Enzyme + PEP*

**B.**

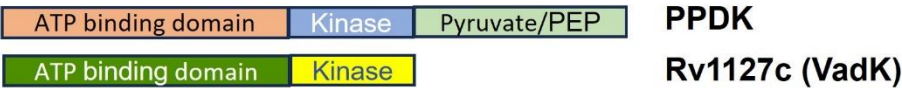

**C.**

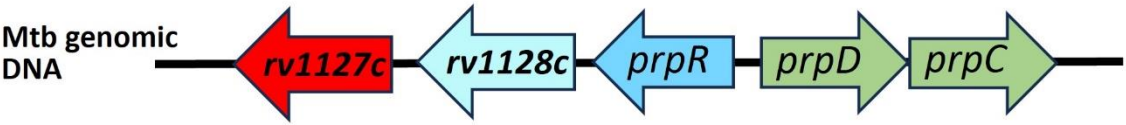

**Figure S1. PPDK and VadK** (A) PPDK catalyses the conversion of pyruvate to phosphoenolpyruvate (PEP) through two independent partial reactions. (B) The gene annotated as *ppdk* (*rv1127c*) in Mtb is truncated and does not encode the PPDK pyruvate/PEP binding domain (C) *rv1127c* (*vadK*) is in close genomic proximity to genes encoding methylcitrate cycle enzymes (*prpC*, *prpD*) and their transcriptional regulator *prpR*.

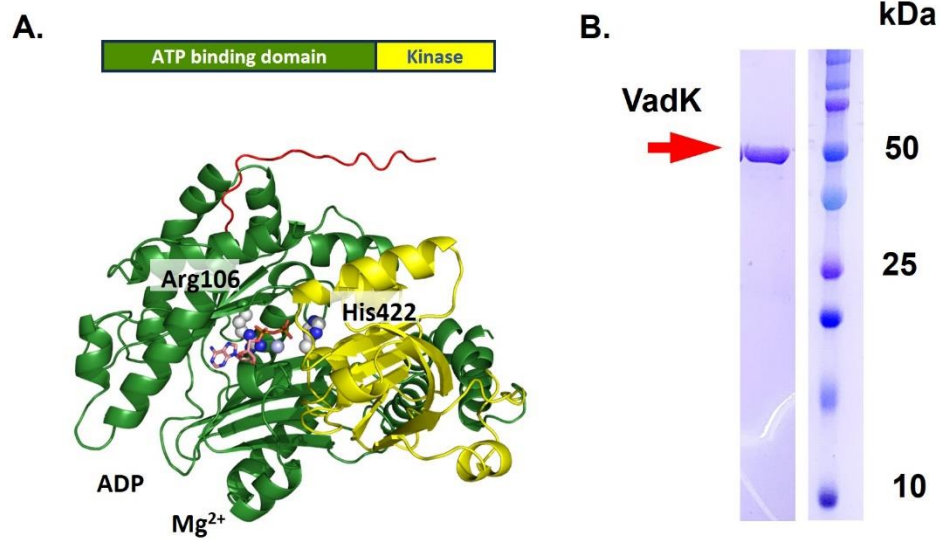

**Figure S2. VadK structure prediction by AlphaFold3 and purified VadK** (A) The N-terminal 1-17 residues, ATP binding and kinase domains colored in red, green and yellow, respectively. ADP and $Mg^{2+}$ are superimposed from *Flaveria trinervia* PPDK (PDB code: 5LU4) and shown in stick (carbons, salmon) and sphere (light blue), respectively. The catalytic histidine, His422, and ATP binding Arg106 are highlighted in spheres (carbons, white). (B) SDS-PAGE gel of purified VadK.

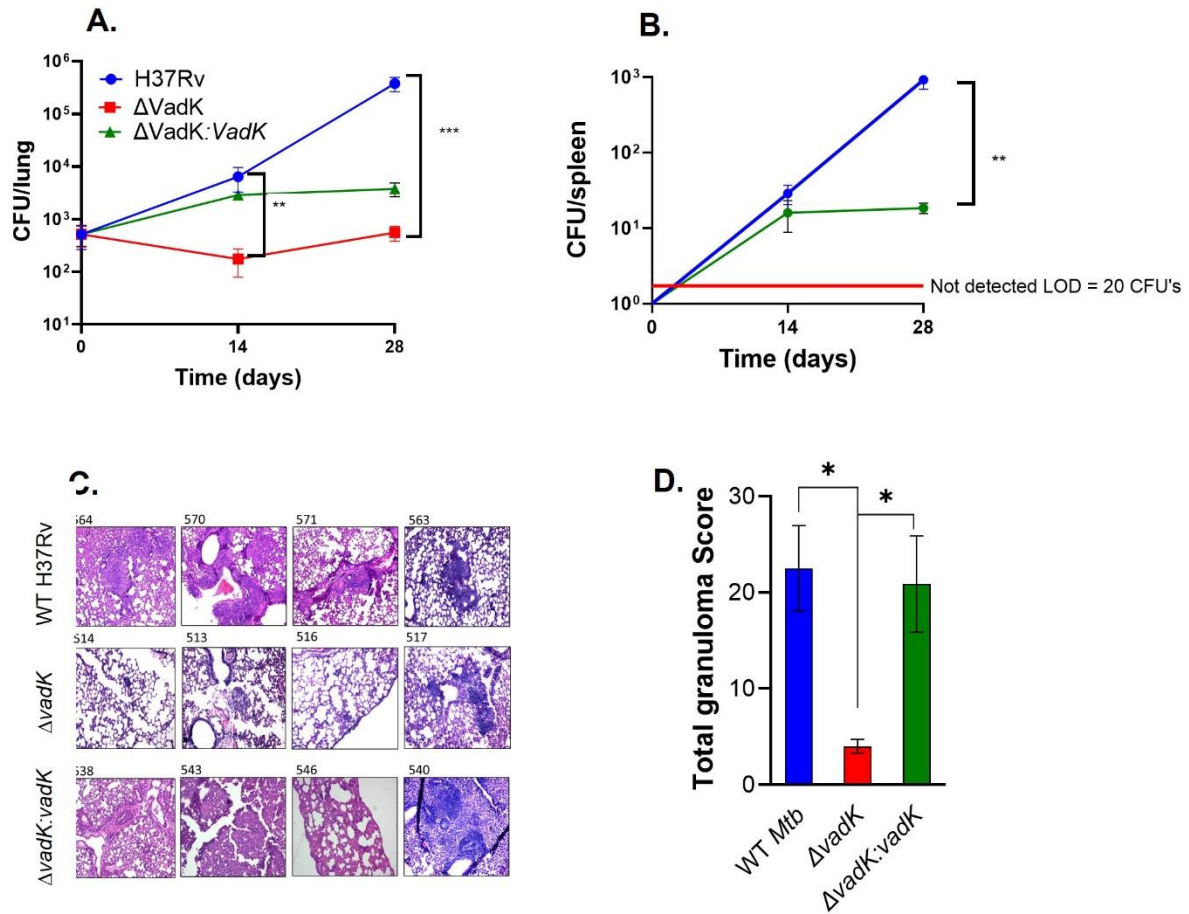

**Figure S3. VadK is required for Mtb to cause tuberculosis in BALB/c mice**
BALB/c mice (n = 6) were aerosol infected with WT (blue), ΔvadK (red) and ΔvadK:vadK (green) Mtb and the bacterial load in the lungs (A) and spleen- ΔvadK not detected (ND) (B) was measured at day 14 and day 28. The limit of detection (LOD) is 20 CFU's. Lung sections were stained with Haematoxylin-and-eosin after 28 days of infection and scored blindly by a pathologist using the method described (doi.org/10.1007/s00281-015-0538-9). The images show HE-stained ($10 \times$ magnification) from individual sections representative of four infected animals (C) and the scores for animals in each group with mean ± standard error (D). The data depicted are means ± SE for each group. $P < 0.001$:\*\*\*, $p < 0.01$:\*\*, $p < 0.05$:\*, unpaired two-tailed t-test with Welch's correction.

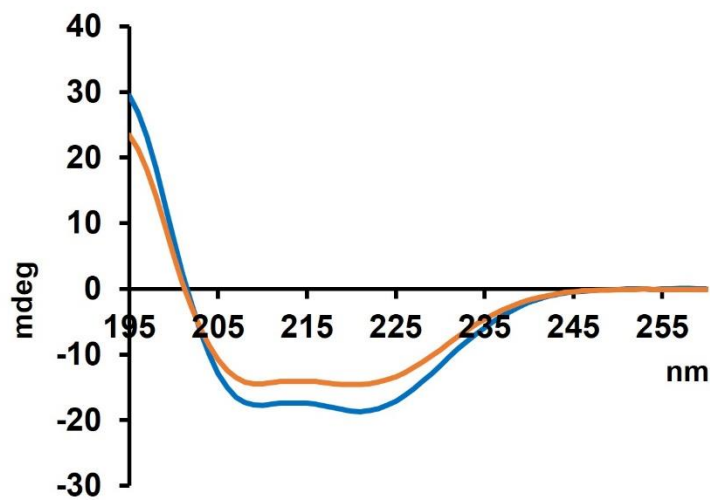

**Figure S4. VadK-H422A is correctly folded.** Circular dichroism spectra of WT VadK (blue line) and VadK-H422A (orange line) are similar, indicating that the mutant has the same secondary structure as WT VadK and is folded correctly.

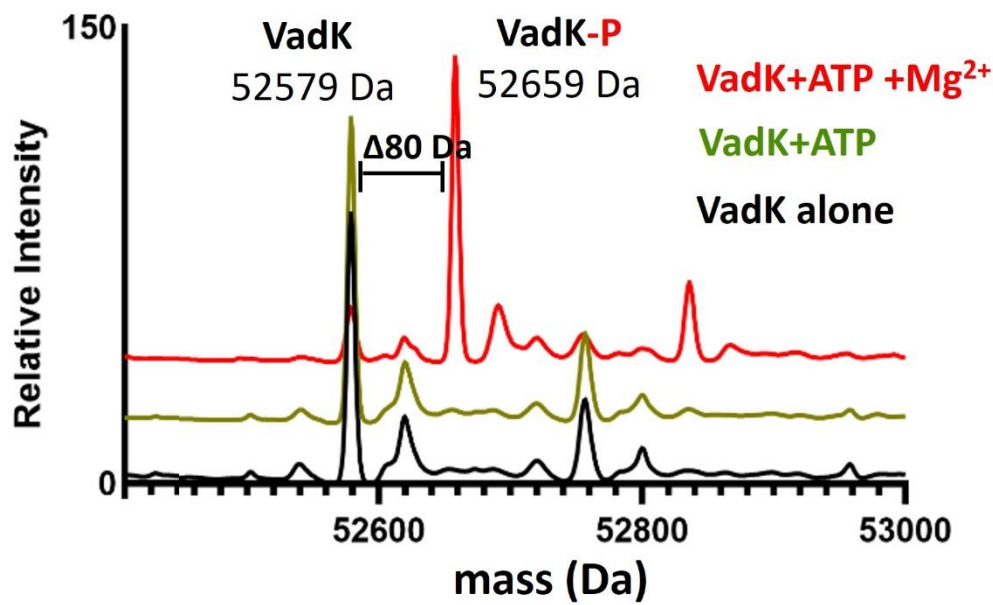

**Figure S5. VadK-His can auto-phosphorylate.** MS of VadK-His alone (black line), with ATP and in the absence (green line) and presence of $Mg^{2+}$ (red line). A phosphorylation event (mass shift 80 Da, designated by VadK-P) only occurs for apo-VadK-His in the presence of ATP and $Mg^{2+}$.

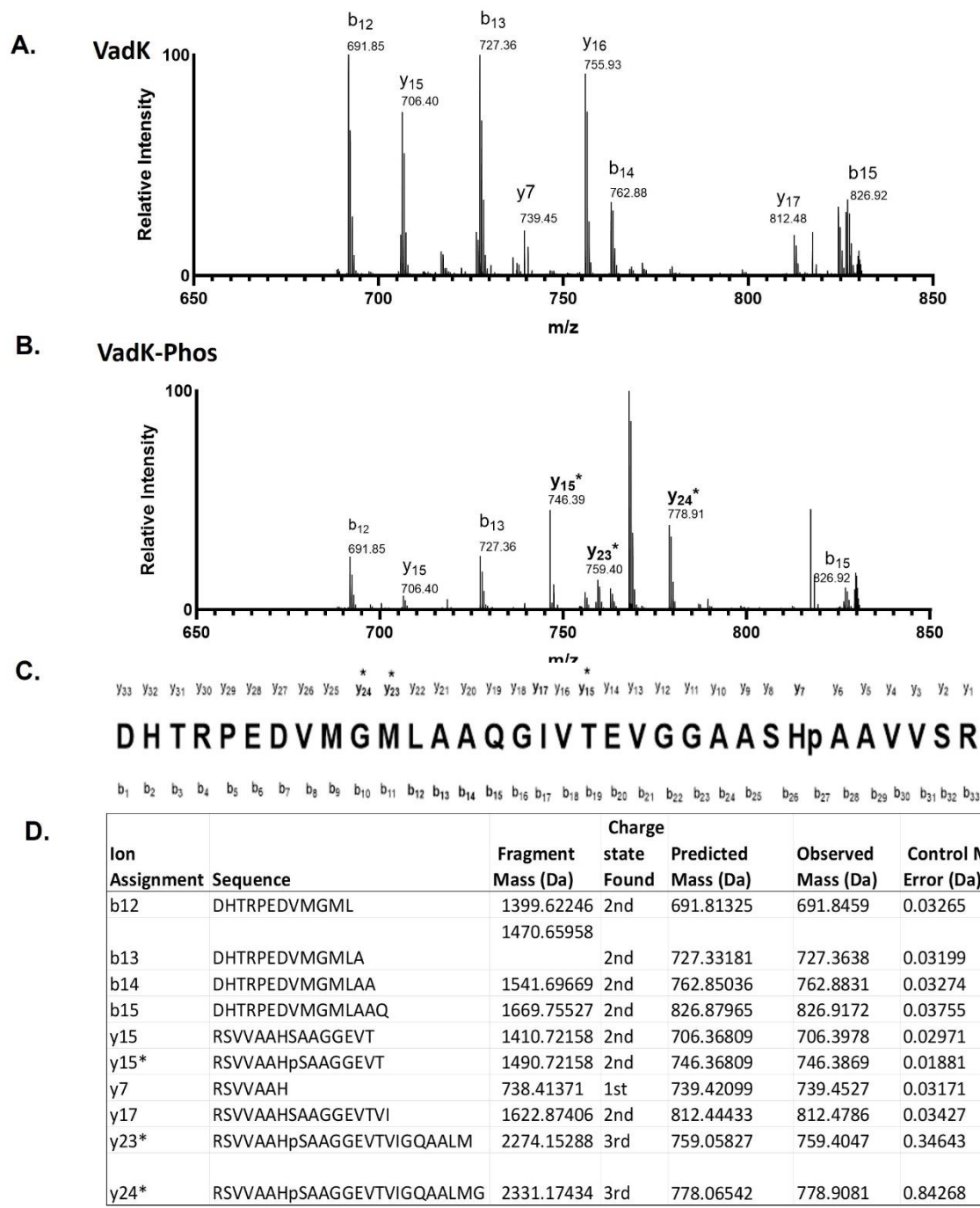

**Figure S6. MS/MS spectra of b/y ion for peptide fragment.** DHTRPEDVMGMLAAQGIVTEVGGAASHpAAVVS R, encompassing amino acid residues 401-433. Comparison of peptide fragment pattern when VadK is (A) untreated and (B) treated with ATP and $Mg^{2+}$. Identification of phosphorylated y ions are labelled with an asterisk and bolded. Notably, the presence of phosphorylated y15 ion suggest that histidine phosphorylation is on H422. (C) peptide sequence denotes order of b and y ions, with p representing phosphorylation modification and identified phosphorylated ion in bold with an asterisk. (D) Table of fragmentation masses which are listed together with the charge state. b and y ions with asterisk indicate phosphorylated ion was identified. Lowercase p in sequence refers to phosphorylation.

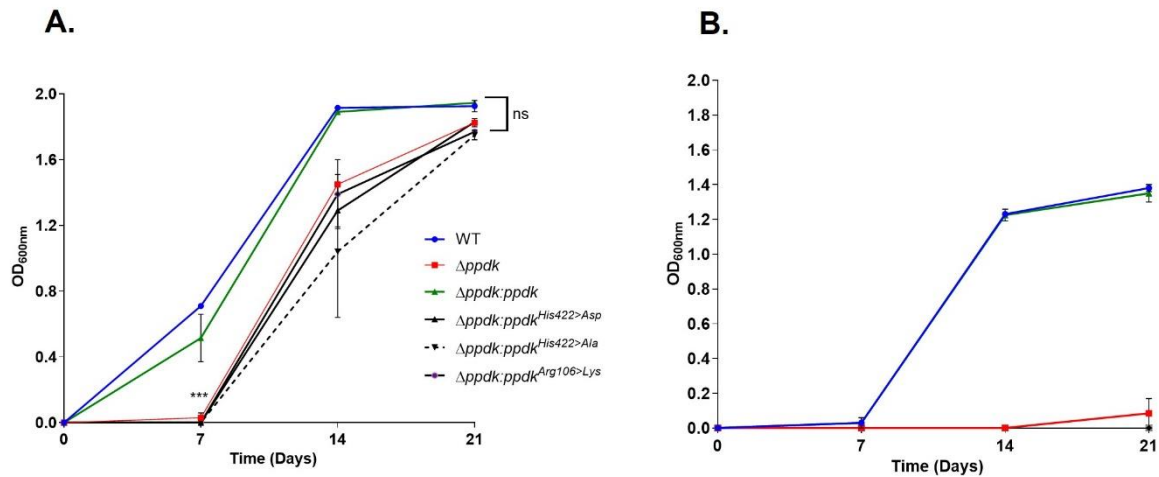

**Figure S7. Strains of Mtb expressing isogenic forms VadK cannot grow in the presence of propionate.** *Mtb*Δ*vadK*, *Mtb*Δ*vadK*:*vadK* and *Mtb* expressing isogenic forms of VadK that is unphosphorylatable because the catalytic histidine has been mutated to alanine (His422>Ala) or is a phosphomimetic because the histidine has been mutated to aspartate (His422>Asp) or unable to bind ATP (Arg106>Lys) were grown in 7H9 media without (A) or with 10 mM propionate (B). Growth was measured by OD (600nm). The data represent the average and SEM of two independent biological replicates. P>0.01:NS, P<0.0001:\*\*\*, two-way ANOVA with Sidak's multiple comparison test.

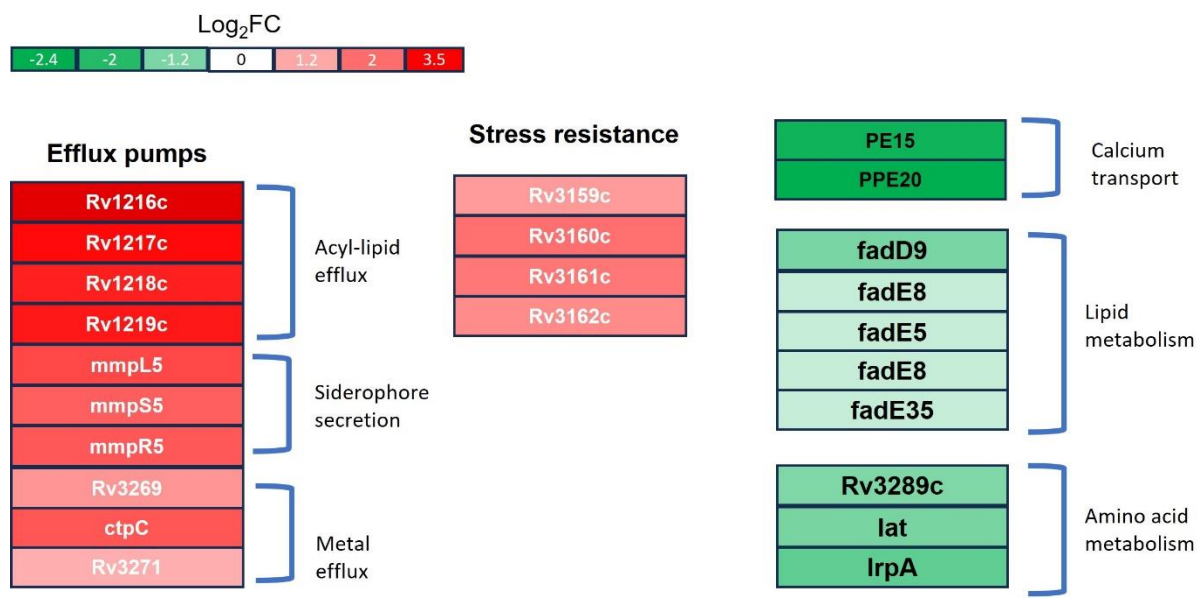

**Figure S8. The absence of VadK is associated with shared transcriptional profile for Mtb growing in cholesterol.** RNA-seq analysis quantifying differentially expressed genes from WT and Δ*vadK* in 7H9 media containing glycerol and 20 mM propionate. Data are displayed as log<sub>2</sub> fold change in gene expression Δ*vadK* vs parental *Mtb*. Data are from two technical replicates from two experiments (\*adjusted q-value ≤ 0.001).

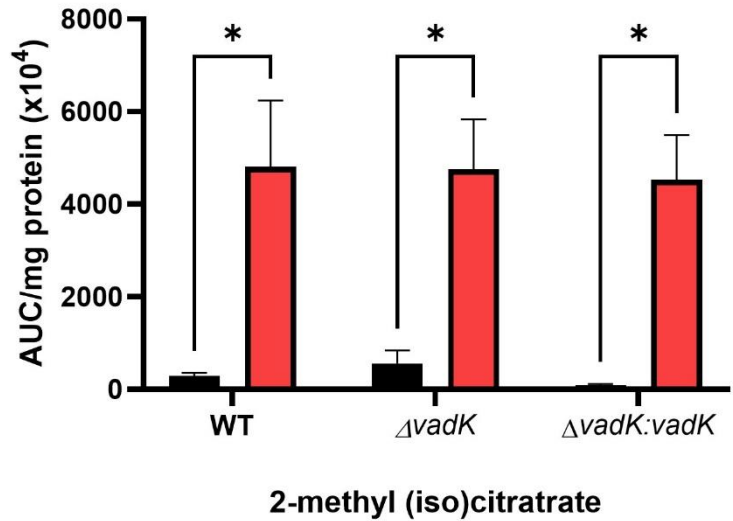

**Figure S9. Abundance of 2-methyl(iso)citrate.** MS measurements of intracellular 2-methyl(iso)citrate from Mtb grown in either Roisin's minimal media with cholesterol (black bars) or 7H9 with 20 mM propionate (red bars) for 48 h. Abundances are shown as normalized AUC (Methods). Mean ± SEM for 3-4 independent biological replicates. \*$P < 0.05$ Student's two-tailed t-test with Welch's correction.

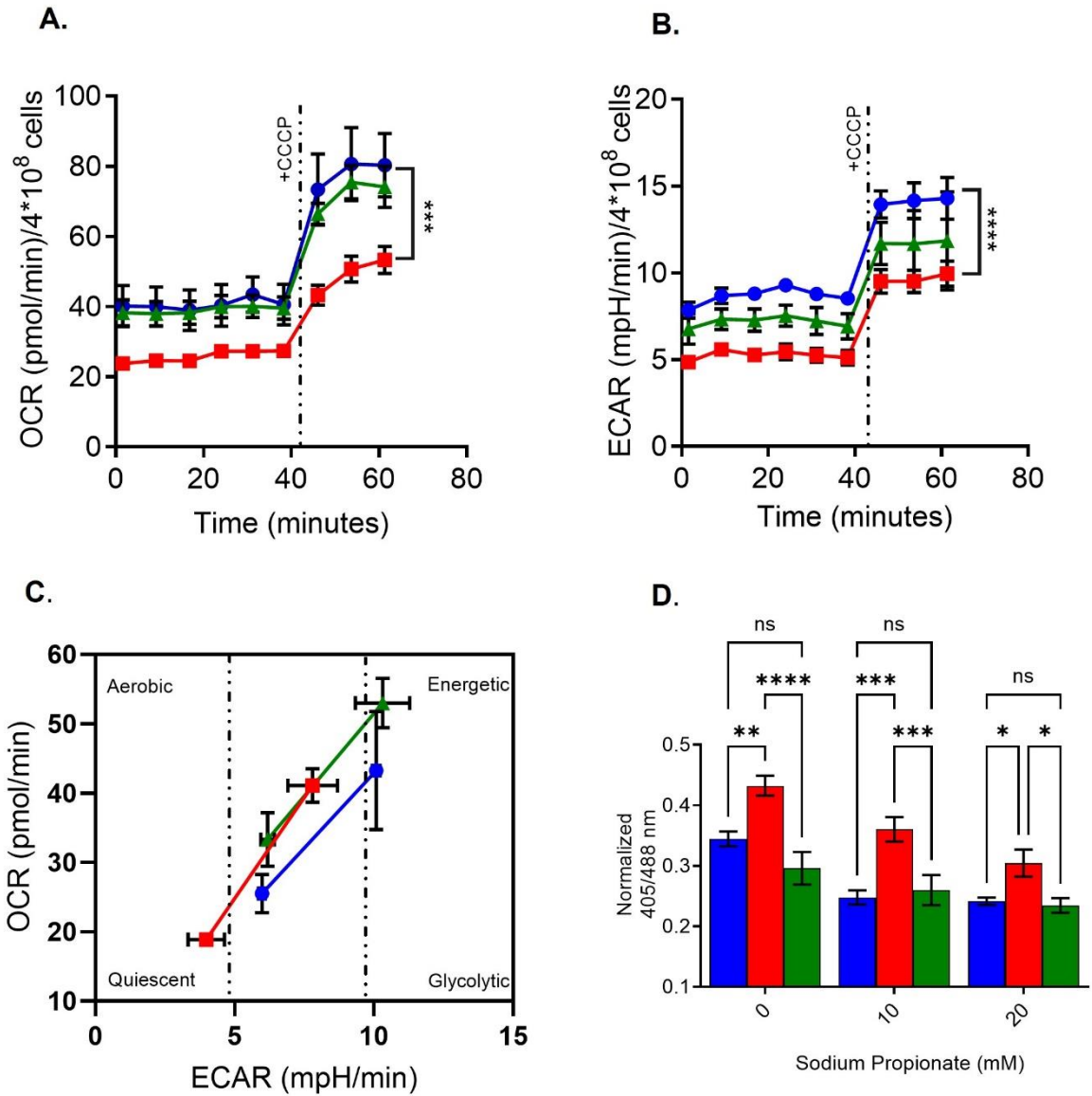

**Figure S10. *VadK* deficient *Mtb* has reduced oxygen consumption rate (OCR), extracellular acidification rate (ECAR) and is more oxidized.** The metabolic potential of WT, $\Delta vadK$, and $\Delta vadK:vadk$ was assessed using the Agilent Seahorse XF Cell Energy Phenotype Test. Basal and stressed energy profiles were generated by measuring (A) oxygen consumption rate (OCR) and (B) extracellular acidification rate (ECAR), before and after treatment with the mitochondrial uncoupler CCCP. (C) The Cell Energy Phenotype Profile indicates that $\Delta vadK$ has low energetics in comparison to WT and complement. OCR and ECAR are normalized to colony-forming units (CFU). Data are presented as means ± SEM. \*$P \leq 0.05$, \*\*$P \leq 0.01$, \*\*\*$P \leq 0.001$, \*\*\*\*$P \leq 0.0001$ calculated by unpaired two-tailed t test. (D) The mycothiol redox potential (EMSH) of WT (blue) and $\Delta vadK$ (red), $\Delta vadK:vadk$ (green) in 7H9 media with glycerol and propionate as indicated was determined by measuring Mrx1-roGFP2 biosensor ratio (405/488 nm) using flow cytometry. The results represent the average and SEM of 3 independent experiments. $P > 0.05$: NS, $P < 0.05$:\*, $P < 0.005$:\*\*, $p < 0.0005$:\*\*\*, $p < 0.00005$:\*\*\*\* two-way ANOVA with Tukey's multiple comparison test.

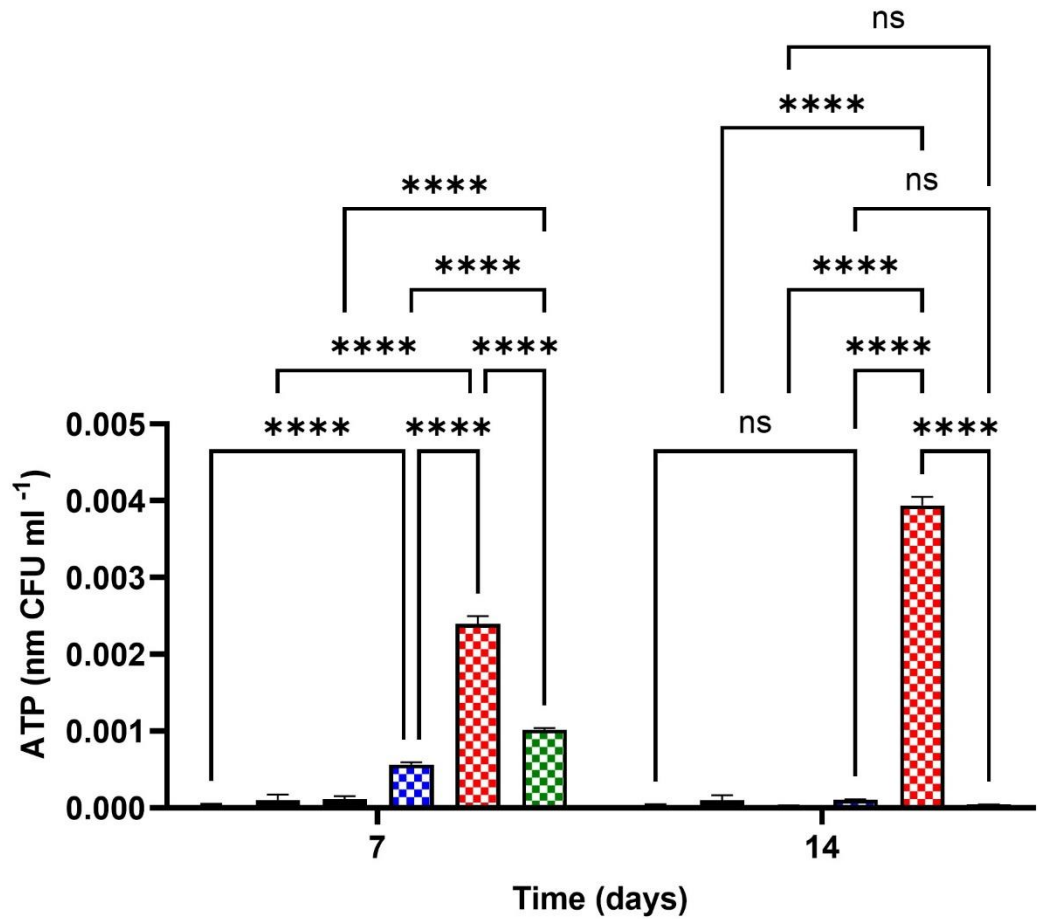

**Figure S11. ATP levels increase in response to propionate in Mtb.** ATP levels in WT (blue bars), $\Delta vadK$ (red bars) and $\Delta vadK:vadK$ (green bars) were grown in 7H9 media without (solid bars) or with 10 mM propionate (checkered bars). ATP was measured (methods) and the results presented are ± SEM for 2-4 independent biological replicates. NS>0.05; \*\*\*\*P < 0.0001; one-way ANOVA with Tukey's multiple-comparison test.

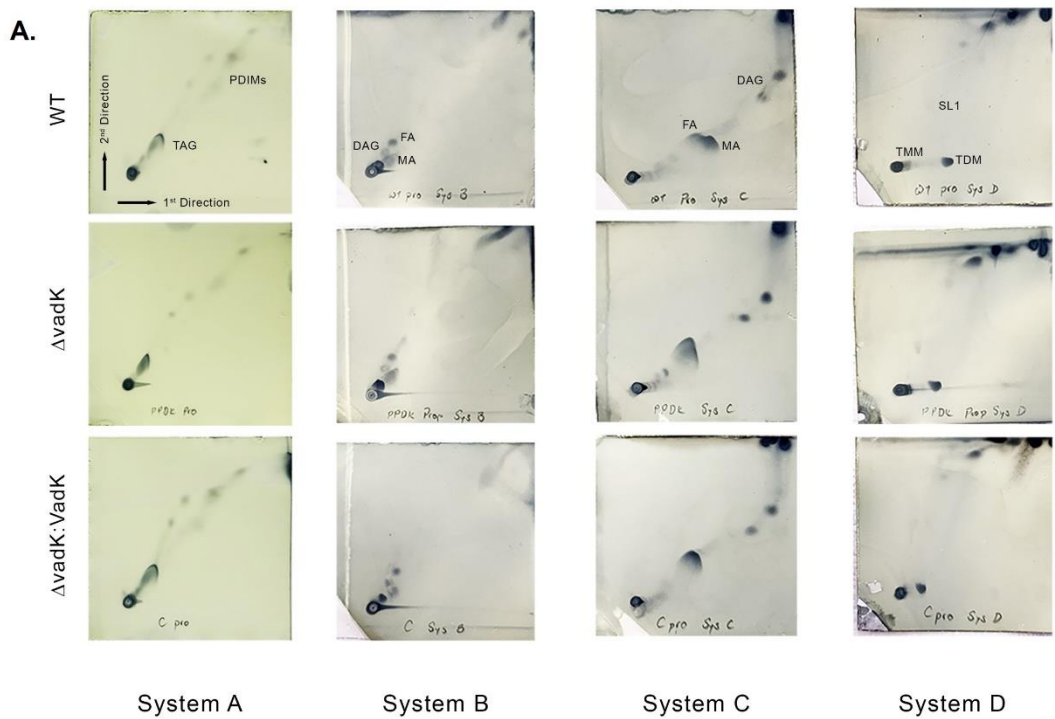

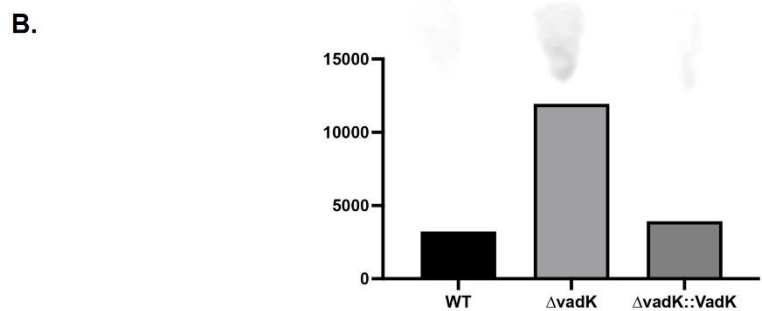

**Figure S12. Lipid composition of VadK deficient Mtb.** (A) 2D TLC analysis of apolar lipid fractions from indicated Mtb strains grown in 7H9 media with 20 mM propionate. 1st and 2nd direction of runs are shown in a representative TLC on the top left-hand corner. TAG; triacylated glycerol, PDIM; phthiocerol dimycocerosate, DAG; diacylated glycerol, TMM, trehalose monomycolate, TDM, trehalose dimycolate; FA, free fatty acid; MA, free mycolic acid; SL1, sulfolipid 1 (B) Bar graph representing densitometric quantification of SL-1 from System D TLCs from different strains. IU, grey pixel intensity unit.
